# APOE Protects Against Severe Infection with *Mycobacterium tuberculosis* by Restraining Production of Neutrophil Extracellular Traps

**DOI:** 10.1101/2024.10.04.616580

**Authors:** Dong Liu, Dat Mai, Ana N. Jahn, Tara A. Murray, John D. Aitchison, Benjamin H. Gern, Kevin B. Urdahl, Alan Aderem, Alan H. Diercks, Elizabeth S. Gold

## Abstract

While neutrophils are the predominant cell type in the lungs of humans with active tuberculosis (TB), they are relatively scarce in the lungs of most strains of mice that are used to study the disease. However, similar to humans, neutrophils account for approximately 45% of CD45+ cells in the lungs of *Apoe^-/-^* mice on a high-cholesterol (HC) diet following infection with *Mycobacterium tuberculosis* (Mtb). We hypothesized that the susceptibility of *Apoe^-/-^* HC mice might arise from an unrestrained feed-forward loop in which production of neutrophil extracellular traps (NETs) stimulates production of type I interferons by pDCs which in turn leads to the recruitment and activation of more neutrophils, and demonstrated that depleting neutrophils, depleting plasmacytoid dendritic cells (pDCs), or blocking type I interferon signaling, improved the outcome of infection. In concordance with these results, we found that Mtb-infected in *Apoe^-/-^*HC mice produce high levels of LTB4 and 12-HETE, two eicosanoids known to act as neutrophil chemoattractants and showed that blocking leukotriene B4 (LTB4) receptor signaling also improved the outcome of tuberculosis. While production of NETs has been associated with severe tuberculosis in other mouse models and in humans, a causative role for NETs in the pathology has not been directly established. We demonstrate that blocking the activation of peptidylarginine deiminase 4 (PAD4), an enzyme critical to NET formation, leads to fewer NETs in the lungs and, strikingly, completely reverses the hypersusceptibility of *Apoe^-/-^*HC mice to tuberculosis.

## INTRODUCTION

Apolipoprotein E (APOE) is a member of a group of lipid binding proteins that plays an important role in lipid transport and metabolism through its interaction with multiple lipoprotein particles including chylomicrons, very-low density lipoprotein (VLDL), and high-density lipoprotein (HDL), and acts as a ligand for their receptor-mediated clearance (*1*). *Apoe* was initially identified as playing a critical role in cardiovascular disease when it was discovered that polymorphisms in APOE lead to familial dysbetalipoproteinemia (*2*). Mice lacking *Apoe* were generated in the 1990s and were shown to develop hypercholesterolemia and atherosclerotic lesions (*3*, *4*) and have served for decades as a major animal model for the study of atherosclerotic vascular disease. One of the main receptors for APOE is the low-density lipoprotein receptor (LDLR) and binding of APOE to the LDLR leads to the clearance of lipoprotein particles from the circulation. Mice lacking *Ldlr* are also hypercholesterolemic and have served as an alternative model for studying atherosclerosis (*5*). While several differences in the models exist, both strains of mice develop similar levels of hypercholesterolemia on high cholesterol diets and, under these conditions, both develop atherosclerotic plaques.

In addition to its role in lipid transport, APOE has been implicated in inflammatory responses (*6*) and has also been shown to play a role in several infectious diseases (*7–12*). An early study found that *Apoe^-/-^* mice on a high-cholesterol (HC) diet are highly susceptible to Mtb-infection and that the susceptibility is increased with increasing hypercholesterolemia (*13*). Surprisingly, *Ldlr^-/-^* mice with similar levels of hypercholesterolemia to *Apoe^-/-^* mice were relatively resistant to Mtb, mounting a timely immune response and demonstrating a similar capacity for controlling the bacteria as wild-type (WT) C57BL/6 (B6) mice (*14*). Because *Apoe^-/-^* mice develop necrotic lesions containing large numbers of neutrophils, similar to those seen in humans with severe tuberculosis, we sought to use this model system to uncover factors leading to severe tuberculosis using *Ldlr^-/-^* mice as a control for any confounding effects of the hypercholesterolemia.

While neutrophils are the most abundant cell type in the lungs of patients with active tuberculosis, comprising 38-86% of cells recovered from cavitary lesions, sputum, or BAL (*15*), they are a relatively small fraction (∼5%) of the responding immune cell population in the most commonly used (and relatively resistant) mouse model of tuberculosis, B6 mice infected with Mtb H37Rv (*16*). In more susceptible strains, excessive neutrophil recruitment (∼10-40% of pulmonary immune cells) has been shown to be detrimental to the control of Mtb, and depleting neutrophils or blocking their recruitment to the lung partially reverses the phenotype (*17–19*). Several factors have been shown to play a role in recruiting neutrophils to the lungs of mice infected with Mtb. Type I interferon is generally considered to be detrimental to control of Mtb infection and several studies have proposed a direct link between excessive type I interferon and neutrophil recruitment (*20–22*). While pDCs represent a relatively small proportion of immune cells in the lung, it has been demonstrated that they are major producers of type I interferon (*23–25*) that can be activated via TLR7 and TLR9 recognition of extracellular DNA (*25*, *26*). In at least one murine model of severe tuberculosis, deletion of *Unc93b*, a chaperone required for TLR7 and 9 function, improved disease outcome as assessed by bacterial burden (*19*). Eicosanoids are lipid mediators of inflammation and several eicosanoids, including LTB4 and 12-HETE, have been demonstrated to be critical mediators of neutrophil swarming and activation state (*27–33*) and, in the context of tuberculosis, it has been shown that 12-HETE promotes excess neutrophil recruitment to the lungs of the highly susceptible *Nos2^-/-^* strain of mice and that this correlates with high bacterial burdens (*17*).

Neutrophils are short-lived innate immune effectors that engage multiple mechanisms to counter invading microbes, including the formation of “neutrophil extracellular traps” (NETs). Formation of NETs is an active process involving citrullination of histones, chromatin decondensation, and extravasation of DNA and associated proteins (*34*). Classically, NET formation depends on activation of the enzyme peptidylarginine deiminase 4 (PAD4) which traffics from the cytoplasm to the nucleus to promote citrullination of histones, and inhibiting PAD4 has been shown to prevent NET formation (*34*). Although PAD4-independent NET formation has been described (*35*), NET formation in Mtb-infected neutrophils is at least partially dependent on PAD4 (*36*).

Several groups have identified a correlation between increased NET formation and susceptibility to tuberculosis in mice (*19*, *20*), and in humans with tuberculosis, NET formation has been associated with necrotic granulomas leading to cavitary lesions (*20*) which are markers of more severe disease leading to prolonged culture positivity, increased risk of relapse and lasting lung damage (*37–39*). However, to the best of our knowledge, no prior publications have shown that blocking NET formation improves the outcome of disease. Importantly, unlike neutrophil depletion, blocking the formation of NETs is not broadly immunosuppressive (*40*, *41*) and in fact, in the case of a model of polymicrobial sepsis, has been shown to offer a survival benefit (*42*).

It has been increasingly appreciated that neutrophils are not a homogenous cell type but can be polarized towards functionally different states. In the context of cancer, it has been proposed that tumor associated neutrophils (TANs) can be polarized into N1 (immunostimulatory) or N2 (immunosuppressive) phenotypes which are considered to be anti- or pro-tumorigenic respectively. N1 neutrophils produce high levels of inflammatory cytokines and other molecules that activate T cells while N2 neutrophils produce multiple matrix-metalloproteases (MMPs) which promote tissue remodeling and angiogenesis (*43–45*).

We observed excessive neutrophil recruitment to the lungs of Mtb-infected *Apoe^-/-^* HC mice as compared to Mtb-infected wild-type (B6) or *Ldlr^-/-^* HC mice and found that depleting neutrophils in *Apoe^-/-^* HC mice partially reversed the excess bacterial burden in these mice. We hypothesized a model where the extreme susceptibility of the *Apoe^-/-^* HC mice was driven by a detrimental feed forward loop in which neutrophils recruited to the lungs produced NETs which in turn activated pDCs to make excess type I interferon which in turn led to further neutrophil recruitment. We tested this hypothesis by blocking each step in the predicted feed-forward loop and demonstrated that, consistent with our model, depleting pDCs or blocking type I interferon signaling partially improved the outcome of Mtb infection. We also found that, at an early time point post-infection (PI), prior to the divergence of bacterial burdens, there are significantly higher levels of the eicosanoids LTB4 and 12-HETE in the serum of Mtb-infected *Apoe^-/-^* HC mice compared to either B6 or *Ldlr^-/-^* HC mice and that blocking the LTB4 receptor improved the outcome of *Apoe^-/-^* HC mice. Most strikingly, we showed that inhibiting PAD4, and thus decreasing NET formation, completely rescued the phenotype of the *Apoe^-/-^* HC mice returning the bacterial burden back to that seen in B6 mice and significantly increasing their survival.

## RESULTS

### *Apoe^-/-^* HC mice are highly susceptible to infection with Mtb

To directly compare the susceptibility of hypercholesterolemic *Apoe^-/-^* and *Ldlr^-/-^* mice, we placed both on a HC diet for two weeks and then infected them via aerosol with approximately 50 CFU of Mtb H37Rv. Both male and female *Apoe^-/-^* HC mice were significantly more susceptible to Mtb than sex and age matched *Ldlr^-/-^*HC mice or B6 HC mice (Figure 1A, Figure S1A) despite broadly similar serum cholesterol profiles of the two knockout strains (Figure 1B, Figure S1B). *Apoe^-/-^*HC mice control Mtb growth with similar efficiency to *Ldlr^-/-^*HC or B6 HC mice for the first 21 days following aerosol challenge but subsequently lose control of the infection and by day 28 have nearly 10-fold higher bacterial burdens (Figure 1C). *Apoe^-/-^* mice on normal chow, while not hypersusceptible to infection with Mtb (Figure 1A), do have an approximately 5-fold higher bacterial burden at day 28 PI than WT mice (Figure S1C).

**Fig. 1.**
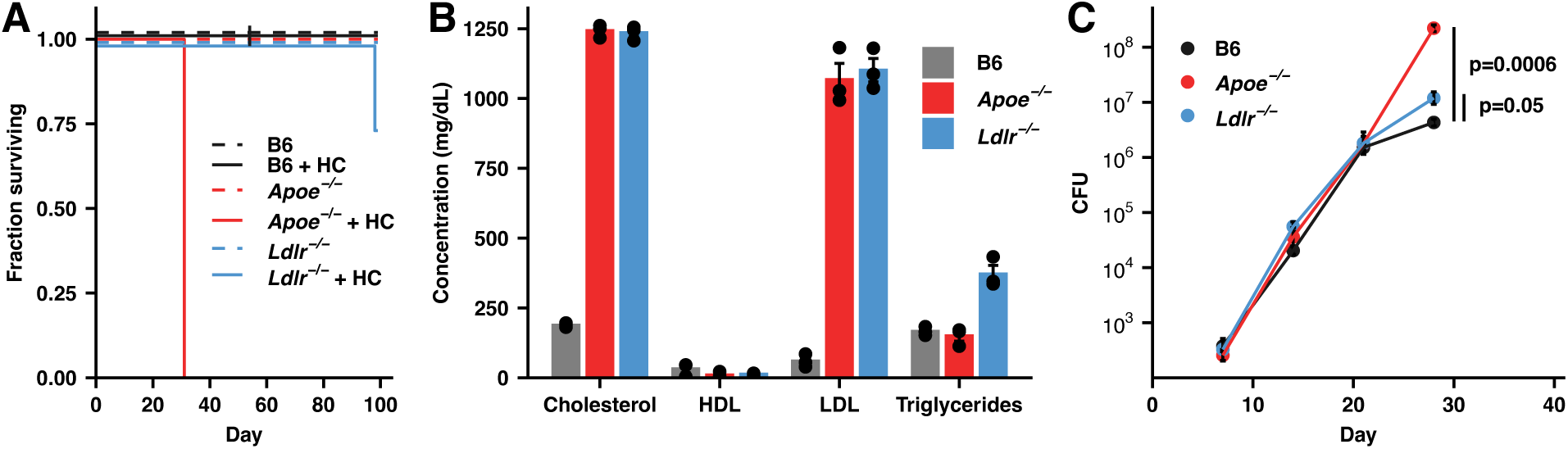
*Apoe^-/-^*HC mice are highly susceptible to infection with Mtb. (**A**) Male mice of the indicated genotypes were fed either normal food or high-cholesterol food for two weeks and then infected with ∼50 CFU Mtb H37Rv and maintained on their pre-infection diet. (n=3 mice/group) (**B**) Serum cholesterol profiles at day 28 following infection of the indicated genotypes of mice fed HC food and infected with Mtb H37Rv as in (A). HDL = high-density lipoproteins, LDL = low-density lipoproteins. (n=3 mice/group) (**C**) Bacterial burden in the lung measured by CFU counting for mice of the indicated genotypes at the indicated time points fed HC food and infected with Mtb H37Rv as in (A). (n=5-7 mice/group) Bars/lines indicate mean; error bars indicate SEM. Significance analysis was performed using the two-sided Student’s t-test allowing for unequal variances (C).

### T cell priming is intact in *Apoe^-/-^* HC mice

In the first publication describing this model, the authors presented evidence, including impaired proliferation of CFSE labeled OT-II OVA-specific T cells stimulated in vivo with OVA-coated beads, suggesting that the extreme susceptibility of *Apoe^-/-^* HC mice to Mtb infection results from defective T cell priming (*13*). However, a recent study demonstrated that APOE deficiency in dendritic cells (DCs) enhances their ability to present antigens to CD4 T cells, resulting in more efficient T cell priming (*46*). We performed a series of experiments to reconcile these disparate findings by evaluating DC and T cell function from *Apoe^-/-^* mice. To test the ability of *Apoe^-/-^* DCs to present model antigens to T cells in vivo, we adoptively co-transferred OVA-specific, CFSE-labeled OT-I and OT-II T cells into either *Apoe^-/-^*, *Ldlr^-/-^* or B6 HC mice. 24hrs later, recipient mice were intranasally (IN) challenged with live recombinant BCG expressing OVA(*47*) (BCG-OVA). Four days later, mice were sacrificed and the expansion of OVA-specific T cells in the mediastinal lymph nodes was measured by CFSE dilution. There was no impairment in the ability of *Apoe^-/-^* DCs to present OVA peptides to either the OT-I or OT-II T cells (Figure 2A). To examine the ability of *Apoe^-/-^*DCs to present Mtb antigens in vivo we adoptively transferred CFSE labeled Mtb-specific (C7) CD4 T cells (*48*), which have been engineered to express T cell receptors specific for the Mtb antigen ESAT6, to *Apoe^-/-^* mice fed either normal (Figure 2B) or HC food (Figure 2C). One day later, the mice were injected intradermally (ID) in the ear with 10^4^ Mtb H37Rv (*49*) and T cell proliferation in the cervical lymph node was examined five days after inoculation. In both cases *Apoe^-/-^, Ldlr^-/-^*, and B6 DCs were equally effective at driving proliferation of exogenous T cells (Figure 2B,C).

**Fig. 2.**
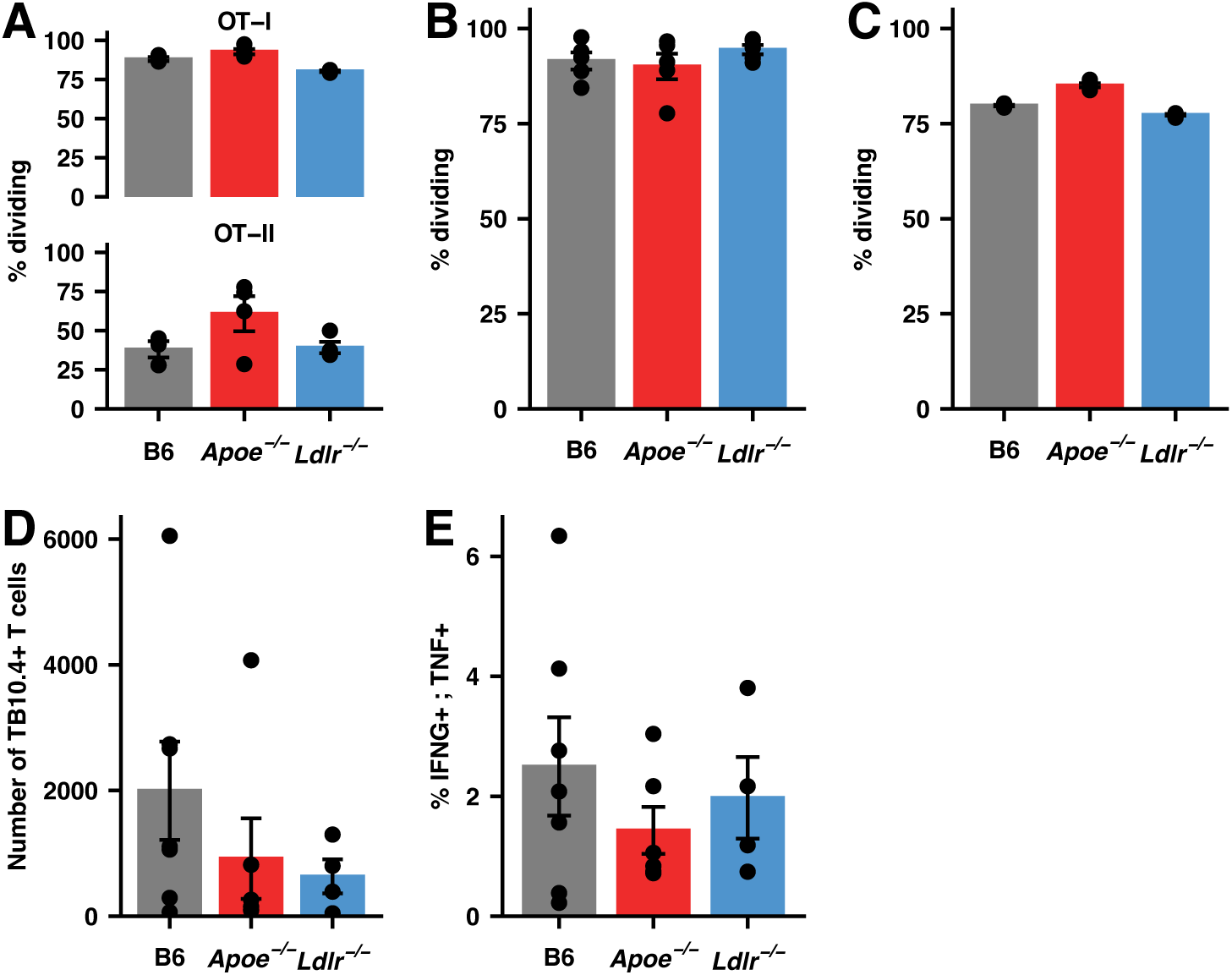
T cell priming is intact in *Apoe^-/-^* HC mice. (**A**) Expansion of CFSE-labeled, CD8 (OT-I) or CD4 (OT-II) T cells specific for Ova peptides as measured by flow cytometry, shown as a percentage of cells dividing in the draining (mediastinal) lymph node, in mice of the indicated genotypes maintained on a HC diet at 4 days following intranasal inoculation with 2×10^8^ CFU BCG-Ova. (n=3-4 mice/group) (**B, C**) Expansion of CFSE-labeled, ESAT-6 specific transgenic CD4+ T cells (C7) as measured by flow cytometry, shown as a percentage of cells dividing in the draining (cervical) lymph node, in mice of the indicated genotypes maintained on a normal (B) or HC (C) diet at 5 days following inoculation with 10,000 CFU Mtb H37Rv in the dermis of the ear. (n=5 mice/group) (**D**) Number of CD8+ TB10.4+ T cells in the lung parenchyma (defined by lack of labeling by an intravenous anti-CD45 antibody (IV-), see Methods) at day 19 following infection with ∼50 CFU Mtb H37Rv in the indicated genotypes of mice maintained on a HC diet. (n=7 mice/group) (**E**) Percentage CD8+ T cells in single-cell suspensions of lung tissue from mice in (D) producing both IFNG and TNF when restimulated with TB10.4 peptides assessed by intracellular staining and flow cytometry. (n=7 mice/group) Bars indicate mean; error bars indicate SEM. Data are representative of 2-4 independent experiments. See Figure S4 for gating strategies.

We also assessed the number of functionally active, Mtb-specific T cells in the lungs of all three genotypes at day 19 PI with ∼50 CFU of H37Rv, a time point where the adaptive immune system has begun to respond to Mtb but that precedes the divergence of bacterial burden (Figure 1C). We found no significant difference in the number of Mtb-specific CD8 (TB10.4 tetramer+) cells in the lungs of *Apoe^-/-^* HC mice when compared to B6 and *Ldlr^-/-^*HC mice (Figure 2D). Furthermore, there was no significant difference in the capacity of these cells to produce IFNG and TNF in response to ex vivo restimulation with the Mtb-specific peptide (Figure 2E).

### The extreme susceptibility of *Apoe^-/-^* HC mice arises from excessive NET formation

We examined the pulmonary cellularity in *Apoe^-/-^*, *Ldlr^-/-^*, and B6 mice on a HC diet over the first 4 weeks of infection and observed that the most striking difference between genotypes was highly elevated levels of neutrophils in *Apoe^-/-^* HC mice (Figure 3A, Figure S2). Taken together with recent studies of other mouse models of extreme susceptibility to Mtb infection which have implicated excessive neutrophil recruitment in the pathology of severe TB disease (*17–19*) and other studies in the literature (*20–26*), these data suggested that the susceptibility of *Apoe^-/-^* HC mice might arise from an unrestrained feed-forward loop in which production of NETs stimulates production of type I interferons by pDCs which in turn leads to the recruitment and activation of more neutrophils (Figure 3B). To test this hypothesis, we disrupted each step in the loop individually and measured the effect on bacterial burden.

**Fig. 3.**
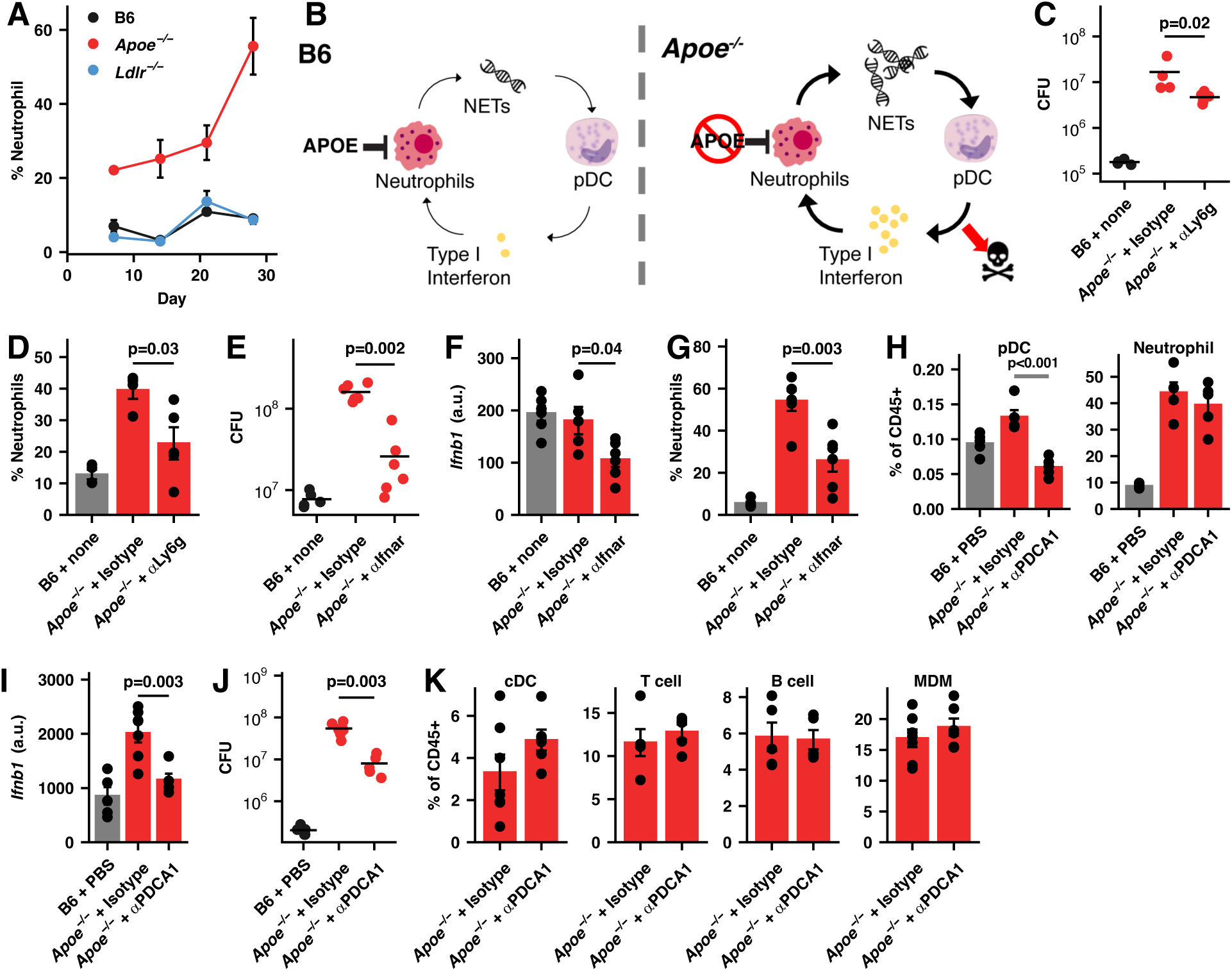
pDC produced Type I interferon contributes to the susceptibility of *Apoe^-/-^* HC mice. (**A**) The kinetics of neutrophil infiltration into the lungs of mice of the indicated genotypes maintained on a HC diet and infected with ∼50 CFU H37Rv, as assessed by flow cytometry, expressed as the percentage of total CD45+ cells in the lung. (n=5-7 mice/group) (**B**) Model for extreme susceptibility of *Apoe^-/-^* HC mice. (**C-K**) *Apoe^-/-^*or B6 mice were placed on a HC diet for two weeks, infected with ∼50 CFU H37Rv via aerosol, and maintained on the diet for the entire experiment. *Neutrophil depletion:* (C) Pulmonary bacterial burden and (D) neutrophil fraction of CD45+ cells at day 24 PI in the indicated treatments. *IFNAR blockade:* (E) Pulmonary bacterial burden, (F) expression of *Ifnb1* mRNA, and (G) neutrophil fractions of CD45+ cells in the lung at day 21 PI in the indicated treatments. *pDC depletion:* (H) Fractions of pDCs and neutrophils among pulmonary CD45+ cells, (I) expression of *Ifnb1* mRNA in the lung, and (J) pulmonary bacterial burden at day 28 PI. (K) Fractions of the conventional DCs (cDCs), T cells, B cells, and monocyte-derived macrophages (MDMs) among pulmonary CD45+ cells for mice the mice in (H). Bars/lines indicate mean; error bars indicate SEM. Data are representative of 2 independent experiments (n=4-7) mice/group) (C-K). Significance analysis was performed using the two-sided Student’s t-test allowing for unequal variances (D, F-H, I) or the Wilcox rank-sum test (C,E,J). See Figure S5 for gating strategies.

As seen in other systems (*17–19*), antibody mediated depletion of neutrophils reduced bacterial burden in the lung (Figure 3C). Notably, in the *Apoe^-/-^* HC model, the number of neutrophils in the lung is higher than in the control mice at a very early time point (day 7 PI) and exceeds 50% of total CD45+ cells in the lung, at day 28 PI (Figure 3A). While we were able to reduce the number of neutrophils, the overall level remained relatively high (Figure 3D).

In some models of severe TB (e.g. C3HeB/FeJ (“Kramnik”) mice (*50*), *Sp140^-/-^* mice (*51*), and mice depleted of GMCSF (*20*)) the excess pathology is largely dependent on dysregulated type I interferon signaling, while in other models (i.e. *Nos2^-/-^* and *Acod^-/-^* mice) the excess bacterial burden appears to be independent of type I interferon and has largely been attributed to dysregulated Il1 signaling (*52*). To establish the role of type I interferon in this system we inhibited the type I interferon receptor (IFNAR) with a blocking antibody. This led to a significant decrease in bacterial burden (Figure 3E) and consistent with our model, blocking IFNAR decreased the total amount of type I interferon and the total number of neutrophils in the lung (Figure 3F,G). pDCs are major producers of type I interferon (*23*, *24*) and antibody mediated depletion of pDCs led to a decrease in *Ifnb1* expression and bacterial burden in *Apoe^-/-^* HC mice without significantly affecting the number of neutrophils in the lung (Figure 3H-J). The depletion antibody, anti-PDCA1, binds to bone marrow stromal cell antigen 2 (BST2), a receptor expressed on several cell types including DCs, mature B cells, and monocytes (*53*), however, we do not measure any significant decrease in these populations (Figure 3K). While we cannot formally rule out a contribution from these other cell types to the decrease in bacterial burden, these measurements, and the fact that expression of *Ifnb1* in the antibody-treated mice returned to WT levels (Figure 3I) suggest that the major effect of the treatment was to deplete pDCs.

To block NET formation, we treated *Apoe^-/-^* HC mice with GSK484 which inhibits production of NETs by inhibiting PAD4, an enzyme required for citrullination of histones during NET formation. Treatment with GSK484 reduced the levels of NETs in the lung as measured by the presence of citrullinated histone H3 (Figure 4A,B) and decreased the expression of *Ifnb1* without decreasing the numbers of neutrophils in the lung (Figure 4C,D). Strikingly, inhibition of NET formation completely rescued the phenotype of the *Apoe^-/-^* mice, returning the bacterial burden to that measured in B6 mice (Figure 4E). To explore the potential clinical efficacy of blocking NET formation, we infected *Apoe^-/-^* HC mice with ∼50 CFU of H37Rv and treated them with GSK484 daily starting at day 7 PI until a pre-specified endpoint of 40 days PI. Treatment with GSK484 significantly decreased mortality compared to controls (Figure 4F). While deletion of *Padi4* has been shown to affect expression of MHC II on tumor associated macrophages (*54*), we did not measure any change in MHCII expression in monocyte-derived macrophages following GSK484 administration (Figure 4G).

**Fig. 4.**
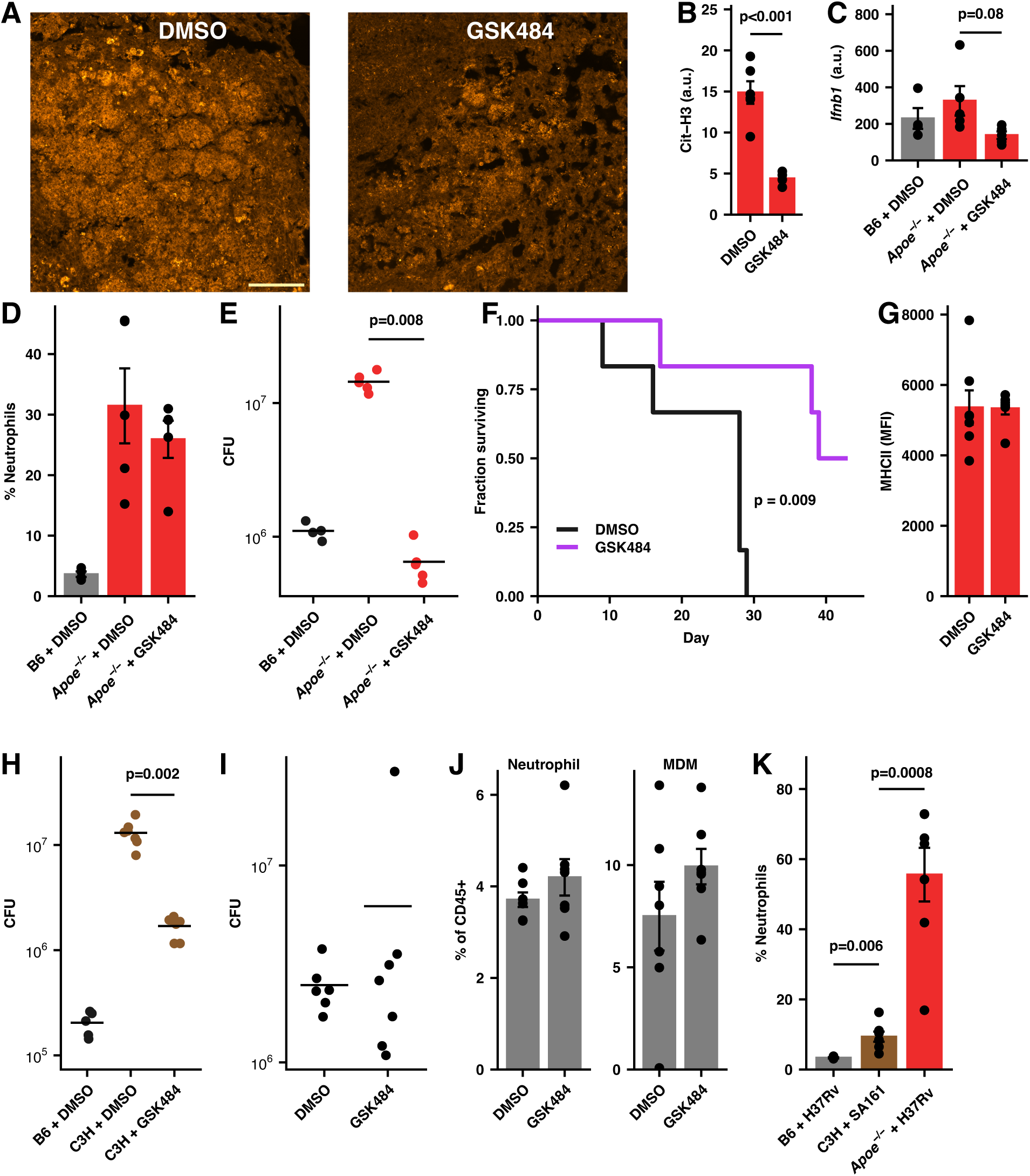
Restraining NET formation protects *Apoe^-/-^* HC mice against severe tuberculosis. (**A**) Representative images of lung sections from Mtb H37Rv infected *Apoe^-/-^*HC mice treated with GSK484 or vehicle daily from days 7-28 PI. Sections were labeled with anti-Cit-H3 antibody (orange) and imaged with confocal microscopy. Scale bar is 100 µm. (**B**) Quantification of the mean fluorescent signal of Cit-H3 labeling for 6 lesions from 3 mice from each condition in (A). (n=6 lesions/group) (**C**) Expression of *Ifnb1* mRNA in the lung, (**D**) fraction of neutrophils among pulmonary CD45+ cells, (**E**) and bacterial burden at day 28 PI in mice treated as indicated. (n=4-5 mice/group) (**F**) *Apoe^-/-^* HC mice were infected and treated as in (A). The fraction of mice surviving to day 40 is plotted. (n=6 mice/group) (**G**) The expression of MHCII on MDM expressed as MFI assessed by flow cytometry from mice treated as in (A). (n=7 mice/group) (**H**) C3H mice were infected with ∼50 CFU Mtb SA161 treated with GSK484 or vehicle daily starting at day 7 PI. Bacterial burden in the lung was measured by CFU at day 28 PI. (n=6-7 mice/group) (**I**) B6 mice were infected with ∼50 CFU Mtb H37Rv and treated with GSK484 or vehicle daily starting at day 7 PI. Bacterial burden in the lung was measured by CFU at day 28 PI. (n=6-7 mice/group) (**J**) The percentage of neutrophils and monocyte-derived macrophages among pulmonary CD45+ cells as measured by flow cytometry at day 28 PI in mice described in (**I**). (**K**) Fraction of pulmonary neutrophils among all CD45+ cells in untreated mice in the infections described in (A), (H), and (I). Bars/lines indicate mean; error bars indicate SEM. Data are representative of two independent experiments (C-E, G). Significance analysis was performed using the two-sided Student’s t-test allowing for unequal variances (B,C,D,G,K), the Wilcox rank-sum test (E,H,I), or the Mantel-Haenszel test (F). See Figure S5 for gating strategies.

To determine the generalizability of these findings, we tested the effect of blocking NET formation in a different mouse strain / bacterial strain combination. C3HeB/FeJ (C3H) mice are highly susceptible to Mtb and the pathology in these mice has been shown to be driven, at least in part, by excess neutrophil recruitment, particularly when infected with the hypervirulent SA161 strain of Mtb (*17*, *55*). Treatment of SA161-infected C3H mice with GSK484 decreased the bacterial burden by approximately 8-fold (Fig 4H). In contrast, blocking PAD4-induced NET formation with GSK484 in B6 mice did not affect the pulmonary bacterial burden or decrease the numbers of neutrophils or monocyte-derived macrophages (Figure 4 I, J). Notably, the effectiveness of blocking NET formation correlates with number of neutrophils in the lungs of the different models tested (Fig 4K) and shows that the treatment is most effective in models with neutrophil levels that most closely match those in humans with active tuberculosis (*15*). Furthermore, these results suggest that GSK484 does not have any significant direct anti-mycobacterial activity under the conditions tested.

### LTB4 receptor signaling contributes to the high bacterial burden in *Apoe^-/-^* HC mice

To search for potential mediators of neutrophil recruitment, we used mass spectrometry to measure the serum concentrations of 44 eicosanoid species (of which 27 were detected in at least one sample) in each genotype following Mtb infection (Figure S3). Although numerous species were strongly upregulated at day 28, when the bacterial burdens in *Apoe^-/-^* HC mice are more than 10-fold higher than those in either *Ldlr^-/-^* HC or B6 HC mice (Figure 1C), both LTB4 and 12-HETE, two eicosanoids that are well-described as neutrophil chemoattractants and activators, were strongly elevated in the serum of *Apoe^-/-^* HC mice prior to the point at which the bacterial burdens diverge between genotypes (Figure 5A). LTB4 and (to a lesser extent) 12-HETE bind to the LTB4 receptor, a pro-inflammatory receptor which is expressed on multiple immune cell types. Blocking LTB4 receptor signaling with CP-105696 in Mtb-infected *Apoe^-/-^* HC mice significantly reduced bacterial burdens compared to controls but, surprisingly, did not affect overall pulmonary neutrophil numbers (Figure 5B,C). Although treatment with CP-105696 has been shown to reduce monocytic infiltration to atherosclerotic lesions (*56*), it did not reduced the levels of pulmonary monocyte-derived macrophages in Mtb-infected *Apoe^-/-^*HC mice (Figure 5D).

**Fig. 5.**
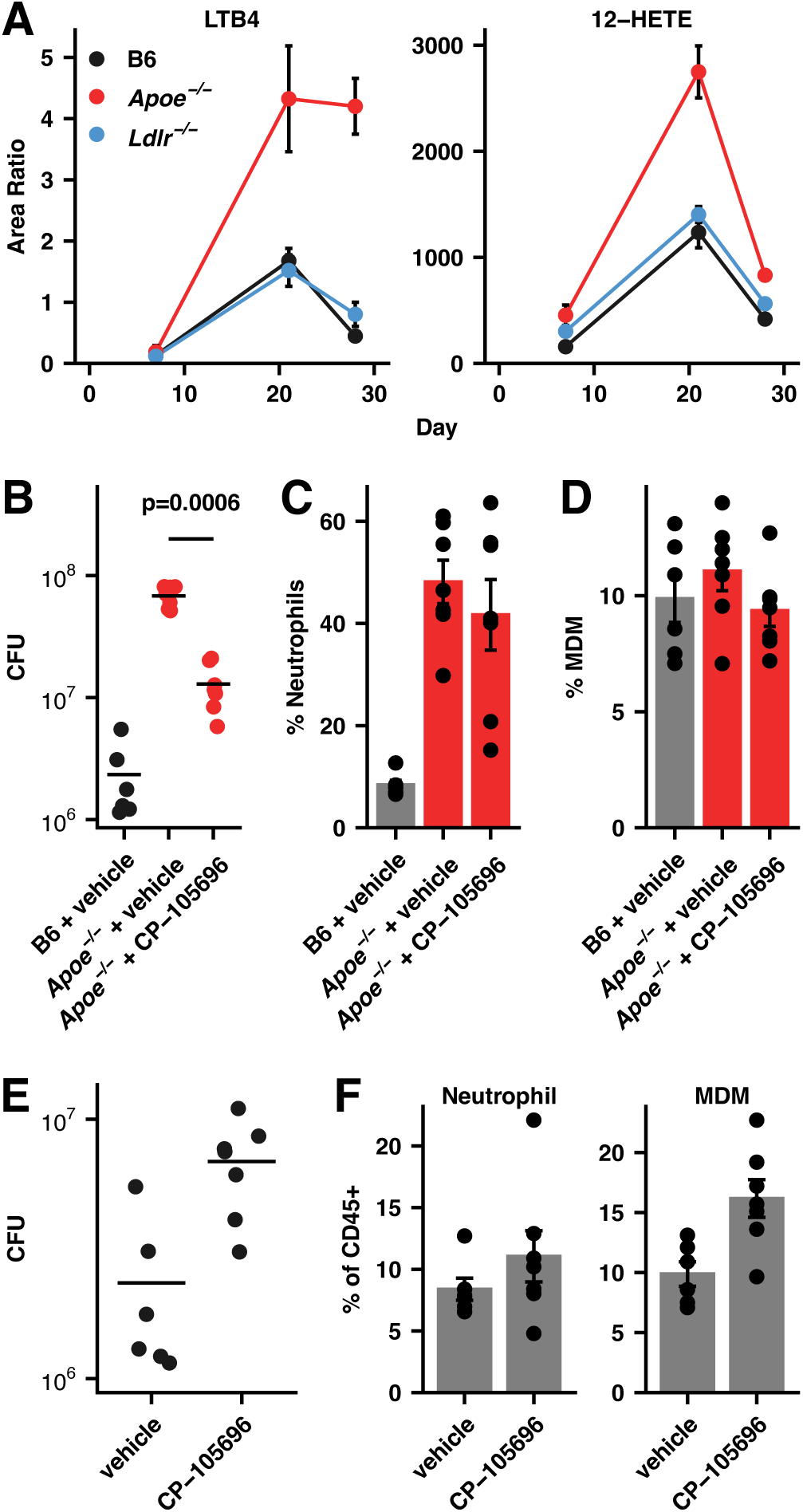
LTB4 and 12-HETE contribute to the hypersusceptibility of Apoe HC mice. (**A**) Mice of the indicated genotypes were infected with ∼50 CFU Mtb H37Rv and maintained on their pre-infection diet. At the indicated time points, levels of LTB4 and 12-HETE in the serum were measured by mass-spectrometry. (n=3 mice/group) (**B**) *Apoe^-/-^* HC mice were infected with ∼50 CFU Mtb H37Rv and left untreated or treated with CP-105696 daily starting at day 7 following infection until day 28. Bacterial burden in the lung was measured at day 28 PI by CFU. (n=6-7 mice/group) (**C,D**) The percentage of neutrophils (C) and monocyte-derived macrophages (D) among pulmonary CD45+ cells as measured by flow cytometry at day 28 PI in mice described in (B). (n=6-7 mice/group) (**E**) B6 mice were infected with ∼50 CFU Mtb H37Rv and left untreated or treated with CP-105696 daily starting at day 7 following infection. Bacterial burden in the lung was measured by CFU at day 28 PI. (n=6-7 mice/group) (**F**) The percentage of neutrophils and monocyte-derived macrophages among pulmonary CD45+ cells as measured by flow cytometry at day 28 PI in mice described in (A). (n=6-7 mice/group) Bars indicate mean; error bars indicate SEM. Significance analysis was performed using the Wilcox rank-sum test (B,E) or the two-sided Student’s t-test allowing for unequal variances (C,D,F). See Figure S5 for gating strategies.

Similar to the results with GSK484, CP-105696 treatment did not improve the outcome of tuberculosis in B6 mice and in fact led to modestly increased bacterial burdens at day 28 without decreasing the numbers of neutrophils or monocyte-derived macrophages in the lungs (Figure 5E,F). These results are concordant with a minimal role for neutrophils in the pathology of Mtb infection in B6 mice (*16*) and suggest that CP-10596 does not have any significant irect anti-mycobacterial activity under the conditions tested.

### Neutrophils in *Apoe^-/-^* HC mice have a distinct polarization state

In several of the experiments described above, the intervention improved the outcome of the mice without affecting the number of neutrophils recruited to the lung. This suggested that the state of the neutrophil when it encounters the Mtb-infected lung may be a critical determinant of disease outcome. To investigate this hypothesis, we examined the transcriptional profiles of *Apoe^-/-^*, *Ldlr^-/-^*, and B6 pulmonary neutrophils at an early time point prior to the divergence in the bacterial burden. We isolated intrapulmonary cells (defined by lack of labelling by an anti-CD45 antibody administered intravenously immediately prior to sacrifice) at day 14 following aerosol challenge with ∼50 CFU of Mtb H37Rv and measured their transcriptomes by single-cell RNA-seq (Figure 6A). When the neutrophil population was isolated and re-clustered, it separated into two distinct populations that were distinguished by expression of numerous genes that correlate with the N1 and N2 phenotypes, as previously described for TANs (*43–45*), including *Tnf*, *Ccl3*, and *Ccl4* (N1) and *Mmp8*, *Mmp9*, and *Ccl6* (N2) (Figure 6B,C). At day 14, the N2 population was significantly larger in the highly susceptible *Apoe^-/-^*HC mice compared to the relatively protected *Ldlr^-/-^* HC mice (Figure 6D).

**Fig. 6.**
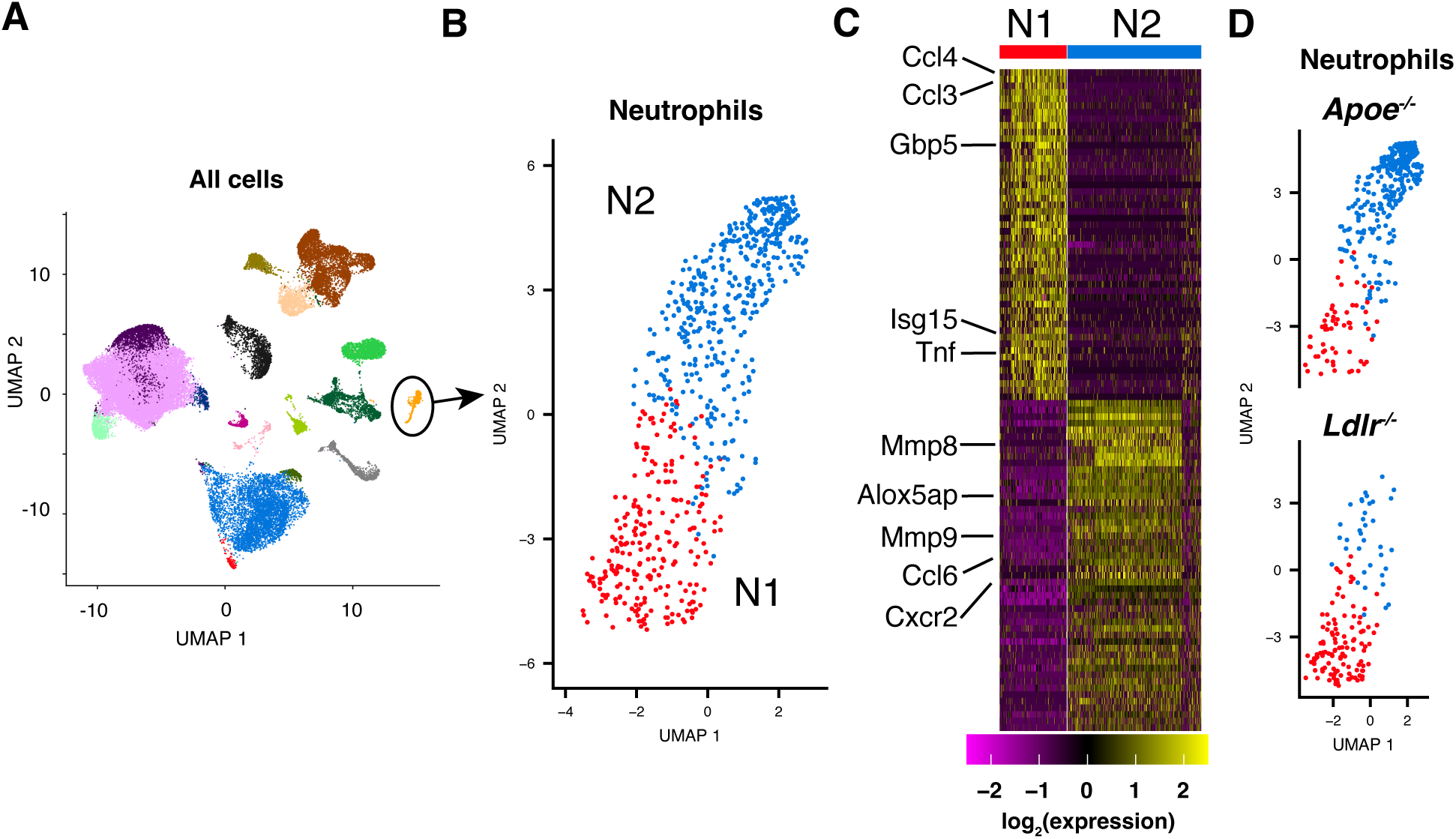
Neutrophils in *Apoe^-/-^*HC mice have a distinct polarization state. (**A**) UMAP plot of expression measurements from single-cell RNA-seq analysis of pulmonary immune cells isolated from B6, *Apoe^-/-^*, and *Ldlr^-/-^* HC mice at day 14 PI with ∼50 CFU Mtb H37Rv (See Methods.). The neutrophil population, identified by comparison with the ImmGen database of transcriptional profiles (https://www.immgen.org) and confirmed by examining expression of *Ly6g* and *S100a8*, is shown in orange and circled. (**B**) UMAP plot of re-clustered expression measurements for the neutrophil population shown in (A). (**C**) Heatmap of row-normalized expression measures for the top 100 genes that distinguish the clusters labeled N1 and N2 in (B). (**D**) UMAP plot of neutrophils from *Apoe^-/-^* and *Ldlr^-/-^* HC mice at day 14 PI. The relative sizes of the N1 and N2 clusters are *Apoe^-/-^* log_2_(N2/N1) = 2.1 ± 1.5 and *Ldlr^-/-^* log_2_(N2/N1) = -1.8 ± 0.3 (mean ± SEM for 3 replicates). See Figure S5 for gating strategy.

This skewing to an N2 phenotype was also observable in *Apoe^-/-^* neutrophils isolated from B6:*Apoe^-/-^*mixed bone-marrow chimeric mice infected with Mtb for 28 days (Figure 7A). Surprisingly, while we measured genotype-specific expression differences between neutrophils in this system, the expression profiles of monocyte-derived macrophages, both infected and uninfected, were quite similar (Figure 7B). A recent paper suggested that APOE, secreted from prostate cancer cells, can bind to TREM2 on neutrophils and drive them towards a senescent phenotype that promotes tumor progression (*57*). Based on the fact that we find that the distinct transcriptional profile of *Apoe*^-/-^ neutrophils is preserved in B6:*Apoe*^-/-^ mixed bone-marrow chimeras on a B6 background (Figure 7A), we do not believe that this mechanism contributes to the pulmonary neutrophil polarization we observe in *Apoe*^-/-^ HC mice.

**Fig. 7.**
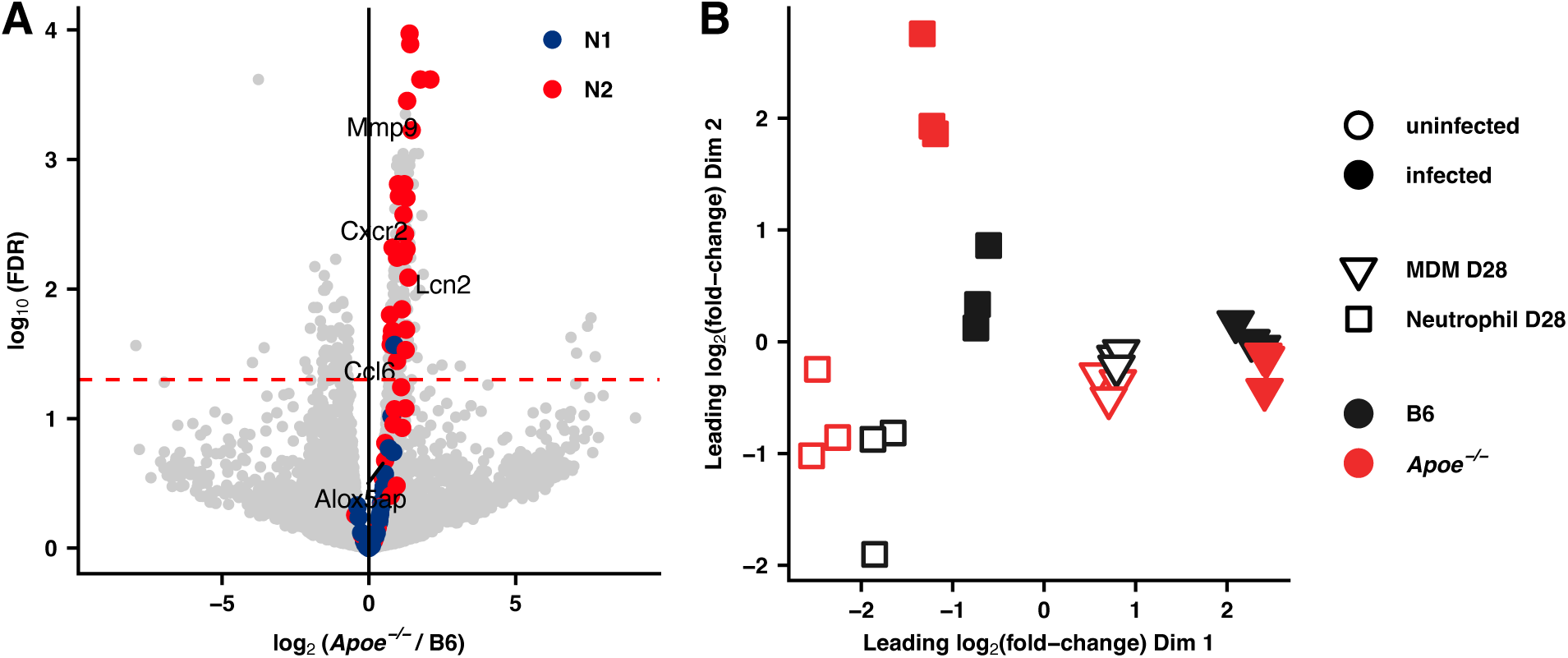
Transcriptional analysis of macrophages and neutrophils isolated from B6:*Apoe^-/-^* mixed bone marrow chimeric mice. (**A**) Volcano plot depicting differential expression between *Apoe^-/-^*and B6 bystander (uninfected) neutrophils isolated from B6:*Apoe^-/-^*mixed bone marrow chimeric mice, maintained on a normal diet, 28 days following infection with ∼50 CFU H37Rv. Genes that are most characteristic of N1 and N2 neutrophils in Mtb-infected mice on a HC diet as determined by single-cell RNA-seq analysis are colored (See Figure 5C). Dashed line indicates FDR=0.05. (n=3 mice/group) (**B**) Multidimensional scaling (MDS) plot^65^ of gene expression in alveolar macrophages (AM), monocyte-derived macrophages (MDM), and neutrophils isolated by cell sorting from B6:*Apoe^-/-^* mixed bone marrow chimeric mice at Day 28 following infection with ∼50 CFU of Mtb H37Rv expressing mCherry^66^. The top 500 genes with the largest standard deviations across samples were used to generate the plot. Distances on the plot represent the leading log2-fold-changes, which are defined as the root-mean-square average of the top largest log2-fold-changes between each pair of samples. (n=3 mice/group) See Figure S5 for gating strategy.

## DISCUSSION

TB is a global health emergency of massive proportions with an annual burden of approximately 10.6 million new cases of active disease and 1.3 million deaths in 2022 (*58*). The recent surge of multi-drug-resistant (MDR) TB (410,000 cases and 160,000 deaths) (*58*), extreme drug resistant TB, and now totally drug resistant TB, emphasizes the need for improved interventions to combat this growing human epidemic. Although standard antibiotic treatment for drug-sensitive TB alleviates active disease for most patients, the rate of recovery and the degree of lung impairment vary significantly. In addition, a minority of patients fail to respond significantly to treatment and continue to harbor detectable bacteria for many months. The specific immune mechanisms responsible for controlling the infection and those that when dysregulated lead to excessive damage remain poorly defined.

We initially hypothesized that a detrimental feed-forward loop involving neutrophil recruitment, production of NETs, activation of pDCs and excess type I interferon production leading to even more neutrophil recruitment was driving the poor outcome seen in *Apoe*^-/-^ HC mice infected with Mtb (Figure 3B). This model was based on reports in the literature that show that both excess neutrophil recruitment and NET production correlate with severe disease (*17*–*20*), that type I interferon is detrimental in Mtb infections and correlates with neutrophil recruitment (*20–22*), and that pDCs are activated to produce type I interferon by the DNA in NETs (*23–25*). While our results are broadly consistent with this original hypothesis, several findings suggest that this model is incomplete. Blocking type I interferon signaling improves disease outcome and decreases neutrophil recruitment to the lungs. However, while depleting pDCs decreases the amount of type I interferon in the lungs (as would be expected) it does not decrease neutrophil recruitment. Furthermore, while blocking NET production dramatically improves disease outcome it does not lead to decreased neutrophil recruitment. Additionally, we identified excess LTB4 and 12-HETE in the serum of the *Apoe*^-/-^ HC mice and found that blocking the LTB4 receptor also improved the outcome of disease but surprisingly did not significantly affect the total number of neutrophils in the lung. These results indicate that this model does not fully describe the complex immune response in vivo and suggests that the state of the neutrophil in the Mtb infected lung, rather than simply the number of neutrophils, is the primary driver of disease outcome.

Patients with severe tuberculosis, characterized by the development of cavitary lesions, are prone to prolonged culture positivity and increased risk of relapse leading to a higher chance of developing MDR TB and to lasting lung damage (*37–39*). We have used the *Apoe*^-/-^ HC mouse which is hypersusceptible to infection with Mtb and which recapitulates several aspects of severe TB in humans, including the development of neutrophil-rich necrotic lesions (*13*), to identify immune mechanisms that are dysregulated during severe disease. Numerous recent studies have highlighted the correlation between high levels of neutrophils in the lung and poor outcome of TB and used depletion strategies to suggest a causal connection (*17–19*). Depletion of neutrophils is unlikely to be a viable clinical treatment and in fact neutropenia is a well-known risk factor for severe disease and death following infection with multiple pathogens. An ideal host directed therapy (HDT) would be highly specific, targeting only those aspects of the immune response that are dysregulated in severe disease while preserving a robust host defense against other infections. However, the specific neutrophil functions that lead to loss of control of Mtb growth or to severe pathology have not been well defined. Our experiments indicate that NET formation is a significant contributor to poor control of Mtb infection during severe disease and suggest that specifically blocking this process can improve outcomes. Unlike neutrophil depletion strategies, blocking the formation of NETs is not broadly immunosuppressive (*40*, *41*) and in fact, in a model of polymicrobial sepsis, has been shown to offer a survival benefit (*42*).

It is increasingly appreciated that mature neutrophils in the periphery can be polarized towards different states. In cancer, polarization of TANs has been shown to play an important role in the immune response: N1 neutrophils are considered inflammatory, express high levels of *Tnf* and restrain tumorigenesis through cytotoxicity and enhancement of anti-tumor responses while N2 neutrophils, which strongly express genes such as *Mmp8* and *Mmp9* are thought to stimulate tumor growth by promoting remodeling of extracellular matrix, enhancing angiogenesis, and inhibiting cytotoxic T-cell responses (*43–45*). While this paradigm has not been well studied in the context of infectious diseases, it is tempting to speculate that in Mtb infection remodeling responses might promote destruction of extracellular matrix which could bias toward necrotic cavity formation and long-term tissue damage. Consistent with this hypothesis we have identified a distinct neutrophil transcriptional profile in *Apoe*^-/-^ mice that are highly susceptible to infection with Mtb which is similar to that described in N2 TANs, while the transcriptional profile of neutrophils from *Ldlr^-/-^* mice that are more resistant to Mtb is similar to that described in N1 TANs. In patients with necrotic granulomatous lesions, adjunctive HDTs that block or modify inflammatory mechanisms that lead to matrix destruction and lung injury, and that enhance antimicrobial drug penetration and action would be particularly useful. Thus, determining whether skewing of neutrophils towards a particular state drives poor outcomes in TB would enable leveraging the ongoing work in cancer to manipulate these states for treating TB and other chronic infectious diseases.

One limitation of this study is that it examines a mouse model, *Apoe*^-/-^, that has extreme hypercholesteremia with lipid levels that are almost never encountered in humans. Because equally hypercholesterolemic *Ldlr^-/-^* mice have essentially identical susceptibility to Mtb infection as wild-type mice, we do not think that hypercholesterolemia alone is driving the extreme TB phenotype in *Apoe*^-/-^ mice. Rather, it appears that the primary causative factor is a massive influx of neutrophils into the lungs of *Apoe*^-/-^ mice (exceeding 50% of pulmonary CD45+ cells) which is not observed in *Ldlr^-/-^*mice (Fig. 4K). Interestingly, the neutrophil levels in Mtb-infected *Apoe*^-/-^ mice are closer than those of other murine models the levels observed in TB patients. In human TB, neutrophils account for ∼86% of cells in sputum, ∼78% of cells in BAL fluid, and ∼39% of cells in the cavities (*15*). In B6 mice infected with H37Rv neutrophils generally only make up ∼5% of the pulmonary CD45+ cells at day 28 PI, and even in the highly susceptible C3H mouse infected with the hypervirulent SA161 strain of Mtb, the neutrophil fraction only reaches ∼10% (Fig. 4K). Therefore, we believe that our findings in *Apoe*^-/-^ mice regarding the role of NETs in the pathology Mtb infection may be particularly relevant to human TB. Based on animal studies, blockade of NETs is postulated to be beneficial in multiple condition including atherosclerotic vascular disease, arthritis, and several types of cancer (*41*, *59–62*). While no PAD4 inhibitors are currently FDA-approved, several are in pre-clinical development by multiple companies (*41*, *62*).

## MATERIALS AND METHODS

### Study Design

The initial objective of this study was to determine the mechanism for the extreme susceptibility of hypercholesterolemic *Apoe*^-/-^ mice to infection with Mtb. We compared the bacterial burdens in *Apoe*^-/-^ HC mice to equally hypercholesterolemic *Ldlr^-/-^* mice over the first 4 weeks following infection with Mtb and determined the point at which the burden’s diverged by CFU analysis of lung homogenates. Previous studies had suggested that the difference in susceptibility might arise from a defect in T cell priming and we tested this hypothesis by examining the expansion of exogenous, antigen-specific T cells and the capacity of endogenous antigen-specific T cells to produce cytokines following Mtb infection. Because these experiments did not indicate a defect in T cell priming or function, we examined the pulmonary cellularity in Mtb-infected *Apoe*^-/-^ HC by flow cytometry and found that *Apoe*^-/-^ HC mice have high levels of neutrophils in the lung.

Taken together with recent studies of other mouse models of extreme susceptibility to, these data suggested that the susceptibility of *Apoe^-/-^*HC mice might arise from an unrestrained feed-forward loop in which production of NETs stimulates production of type I interferons by pDCs which in turn leads to the recruitment and activation of more neutrophils (Figure 3B). To test this hypothesis, we disrupted each step in the loop individually (depleting neutrophils or pDCs, blocking IFNAR signaling, and blocking NET formation) and measured the effect on bacterial burden. To identify mechanisms that lead to high levels of neutrophils and NET formation in *Apoe^-/-^*HC mice, we measured the circulating levels of a panel of eicosanoid species by mass-spectrometry and found that these mice have elevated levels of LTB4 and 12-HETE. To examine the role of these species in mediating pathology, we blocked their common receptor pharmacologically during Mtb infection and examined the effect on bacterial burden. To more comprehensively compare the state of neutrophils in the lung between *Apoe^-/-^* HC mice and controls, we measured their transcriptomes by single-cell and bulk RNA-seq analysis.

Group sizes were determined by analyzing the variance in similar previous experiments or in pilot studies and no statistical methods were used to predetermine group sizes. The time point to stop data collection in the experiment testing the efficacy of GSK484 was predefined based on our prior observations of the time to death in untreated *Apoe^-/-^* HC mice. No data were excluded. Experiments were replicated as indicated in the figure captions. Mice and samples were analyzed unblinded; however, the order of sample groups was generally random during processing and analysis such that the experimentalist could not easily identify the groups.

### Mice

WT (C57BL6/J, JAX:000664, RRID:IMSR_JAX:000664), *Apoe^-/-^*(B6.129P2-Apoe^tm1unc/J^, JAX:002052, RRID:IMSR_JAX:002052), *Ldlr^-/-^*(B6.129S7-Ldlr^tm1Her/J^, JAX:002207, RRID:IMSR_JAX:002207), WT CD45.1 (B6.SJL-Ptprc^a^ Pepc^b^/BoyJ, JAX #002014, RRID:IMSR_JAX:002014), C3H (C3HeB/FeJ, JAX#000658), OT-I (C57BlV6-Tg(TcraTcrb)1100Mjb/J, JAX#003831, RRID:IMSR_JAX:003831), OT-II (B6.Cg-Tg(TcraTcrb)425Cbn/J, JAX#004194, RRID:IMSR_JAX:004194), and ESAT-6 TCR Tg (C7) (JAX035728, RRID:IMSR_JAX:035728) strains of *Mus musculus* were obtained from the Jackson Laboratories (Bar Harbor, ME). OT-I and OT-II mice were crossed onto the CD45.1 mice. All KO mouse experiments used only homozygous animals. All mice were housed in group housing not exceeding 5 animals per cage and maintained in specific pathogen–free conditions at the Seattle Children’s Research Institute (SCRI). Mice were maintained on standard chow (PicoLab Rodent Diet 20, LabDiet). In experiments where mice were fed a HC diet the animals were switched to diet D12109C (Research Diets) 14 days prior to infection and maintained on this diet throughout the remainder of the experiment. Healthy 8- to 14-week-old mice without any previous procedure history were used for all experiments and age and sex matched within each experiment. Our study examined male and female animals, and the findings were similar for both sexes.

### T cell priming assays

OVA-specific CD4+ or CD8+ T cells or TB-specific CD4+ T cells were prepared from spleen and lymph nodes of OT-II, OT-I, or C7 TCR transgenic mice by negative selection using the CD4+ or CD8+ T Cell Isolation Kit (Miltenyi Biotec., #130-104-454, #130-104-075) according to the manufacturer’s instructions. For T cell proliferation assays, purified T cells were labeled with 2 μM CFSE (ThermoFisher, #C34554) before transfer. 10^6^ purified OT-II CD4+, OT-I CD8+, or C7 CD4+ T cells were adoptively transferred into mice by retro-orbital injection. 24 hours later, mice were infected either IN (for BCG-OVA) or ID (for H37Rv) with the indicated doses of live BCG-OVA or H37Rv. Draining lymph nodes were harvested 4 days (mLNs for BCG-OVA) or 5 days (cLNs for H37Rv) PI, and single-cell suspensions were prepared, stained and fixed, and then analyzed on a LSRII or A5 flow cytometer (BD Bioscience).

### T cell function assays

Single-cell suspensions were made from murine lung and were stimulated with TB10.4 (IMYNYPAM) peptide (5 μg/mL final concentration) for 4-6 hours in complete RPMI 1640 media in the presence of 1 μg/mL anti-CD28, anti-CD49d, and Brefeldin A (10 μg/mL) at 37 °C with 5% CO_2_. Cells were washed and surface stained for 30 minutes in the dark at 4 °C, then fixed and permeabilized using BD Cytofix/Cytoperm™ Fixation/Permeabilization kit (Cat # 555028) for intracellular cytokine staining.

### Mouse Mtb aerosol infection

For standard-dose (∼50 CFU) infections, mice were enclosed in an aerosol infection chamber (Glas-Col) and frozen stocks of bacteria were thawed, diluted 1:75 in 0.01% Tween-80 in water, and placed inside the associated nebulizer. To determine the infectious dose, three mice in each infection were sacrificed after the aerosolization was complete. The whole lung was homogenized in 0.05% Tween-80 in PBS with a gentleMACS Tissue Dissociator (Miltenyi Biotec) and serial dilutions were plated onto 7H10 plates for CFU enumeration, as described previously(*63*). All infections used the H37Rv strain of Mtb unless otherwise indicated. The SA161 strain of Mtb was provided by Ian Orme (Colorado State University).

#### Neutrophil depletion

Mice were fed a HC diet for 14 days prior to infection. Mice received either 200 μg of anti-Ly6g antibody (Anti-mouse Ly6G, BioXCell, Cat #BP0075-1, RRID:AB_1107721) or of isotype control (Rat IgG2a, BioXCell, Cat#BP0089, RRID: AB_1107769) via IP injection starting 2 days prior to infection and then every 3 days thereafter for the duration of the experiment.

#### pDC Depletion

Mice were fed a HC diet for 14 days prior to infection. At days -3 and -1 prior to infection and then every 5 days thereafter throughout the experiment, mice received 0.25 mg of anti-PDCA1 antibody (Anti-mouse CD317, BioXCell, Cat #BE0311, RRID:AB_2736991) or isotype control antibody (Rat IgG2b, BioXCell, Cat#BE0090, RRID:AB_1107780) via IP injection.

#### Blocking IFNAR1

Mice were fed a HC diet for 14 days prior to infection. Mice received either 0.5 mg of anti-IFNAR1 antibody (Leinco Technologies, Clone MAR1-5A3, RRID:AB_2830518) or of isotype control (Leinco Technologies, Clone MAR1-5A3, RRID:AB_2830518) via IP injection starting 2 days prior to infection and then every 3 days thereafter for the duration of the experiment.

#### Inhibiting PAD4

Mice were fed a HC diet for 14 days prior to infection. Mice received an IP injection of 0.2 mg of GSK484 (MedChem Express, HY-100514) or PBS with 4% DMSO daily starting 7 days post-infection for the duration of the experiment.

#### Blocking the LTB4R

Mice were fed a HC diet for 14 days prior to infection. Mice received either 1 mg daily of CP-105696 (MedChem Express, HY-19193) in the solvent (10% DMSO, 40% PEG 300, 5% Tween-80 and 45% Saline) or solvent only via oral gavage starting at day 7 PI and daily thereafter for the duration of the experiment.

### Bone marrow transplantation

Bone marrow was harvested by flushing the femurs of the donor mice. B6 CD45.1 mice (B6.SJL-Ptprc^a^ Pepc^b^/BoyJ) were irradiated with two doses of 500 rads using an X-Rad 320 irradiator, then reconstituted with 10^6^ bone marrow cells as, 10^6^ cells from 1:1 mix of *Apoe^-/-^*(CD45.2) and B6 CD451/2 (C57BL/6 x B6.SJL-Ptprc^a^ Pepc^b^/BoyJ) bone marrow. Mice were allowed to recover for 8 weeks and then placed on HC diet for 14 days prior to infection.

Engraftment was confirmed by flow cytometry.

### Cell sorting and flow cytometry

Samples for flow cytometry and cell sorting were prepared as described previously(*63*). Significant details are presented here.

#### Isolation of single-cell suspensions from lung

For T cell experiments, at the indicated times post-infection, mice were anesthetized with isoflurane and administered 1 μg APC-labeled anti-CD45 antibody intravenously to distinguish cells in the circulation (IV+) from those in the lung parenchyma (IV-). Five minutes later, mice were euthanized by CO_2_ asphyxiation, lungs harvested in HEPES buffer containing Liberase Blendzyme 3 (70 μg/mL; Roche, #05401020001) and DNaseI (30 μg/ml; Sigma-Aldrich, #10104159001), and lightly homogenized using a gentleMacs dissociator (Miltenyi Biotec). The lightly homogenized lungs were then incubated for 30 min at 37 °C and then homogenized a second time using the gentleMacs. The homogenates were filtered through a 70 μm cell strainer, pelleted for RBC lysis with ACK lysing buffer (ThermoFisher, #A1049201), and resuspended in FACS buffer (PBS containing 2.5% FBS).

#### Flow cytometry Analysis and Antibodies

For surface staining, cells were suspended in 1X PBS (pH 7.4) containing 0.01% NaN_3_ and 1% fetal bovine serum and blocked with anti-CD16/32 (2.4G2, BD Bioscience), then labeled at 4 °C for 30 minutes in the dark. For intracellular cytokine detection, cells were surface stained and fixed, and then permeabilized. Cell viability was assessed using Live/Dead fixable Aqua or Blue dye (ThermoFisher, #L34966, #L23105). Stained cells were analyzed on a BD LSR II or A5 flow cytometer (BD Bioscience). Samples for flow cytometry were fixed in 2% paraformaldehyde solution in PBS and analyzed using a LSRII or A5 flow cytometer (BD Biosciences) and FlowJo software (Tree Star, Inc.).

The following reagents were used for flow cytometry analysis:

BST2: PE anti-mouse CD317 (BST2, PDCA-1) Antibody (927) (BioLegend, Cat # 127010, RRID:AB_1953285)
CD4: BD Horizon™ BUV496 Rat Anti-Mouse CD4 (GK1.5) (BD Biosciences, Cat # 612952, RRID:AB_2813886)
CD8a: BD Horizon™ BUV395 Rat Anti-Mouse CD8a (53-6.7) (BD Biosciences, Cat # 563786, RRID:AB_2732919)
CD11b: Brilliant Violet 570™ anti-mouse/human CD11b Antibody (M1/70) (BioLegend, Cat # 101233, RRID:AB_10896949)
CD11c: Brilliant Violet 605™ anti-mouse CD11c Antibody (N418) (BioLegend, Cat # 117333, RRID:AB_11204262)
CD11c: APC/Fire™ 750 anti-mouse CD11c Antibody (N418) (BioLegend, Cat # 117352, RRID_AB_2572124)
CD16/32: TruStain FcX™ (anti-mouse CD16/32) Antibody (BioLegend, Cat # 101320, RRID:AB_1574973)
CD19: BD OptiBuild™ BUV563 Rat Anti-Mouse CD19 (1D3) (BD Biosciences, Cat # 749028, RRID:AB_2873425)
CD45: FITC anti-mouse CD45 Antibody (30-F11) (BioLegend, Cat # 103108, RRID: AB_312973)
CD45: PerCP/Cyanine5.5 anti-mouse/human CD45R/B220 Antibody (RA3-6B2) (BioLegend, Cat # 103236, RRID:AB_893354)
CD45: APC anti-mouse CD45 Antibody (30-F11) (BioLegend, Cat # 103112, RRID:AB_312977)
CD64: PE/Cyanine7 anti-mouse CD64 (FcγRI) Antibody (X54-5/7.1) (BioLegendCat # 139314, RRID:AB_2563904)
Ly6c: BD OptiBuild™ BUV805 Rat Anti-Mouse Ly-6C (HK1.4.rMAb) (BD Biosciences, Cat # 755202, RRID:AB_11204262)
Ly6c: Brilliant Violet 785™ anti-mouse Ly-6C Antibody (HK1.4) (BioLegend, Cat # 128041, RRID:AB_2565852)
Ly6g: Brilliant Violet 711™ anti-mouse Ly-6G Antibody (1A8) (BioLegend, Cat # 127643, RRID:AB_2565971)
MHCII: BD OptiBuild™ BUV615 Rat Anti-Mouse I-A/I-E (M5/114.15.2) (BD Biosciences, Cat # 751570, RRID:AB_2875565)
MHCII: Brilliant Violet 650™ anti-mouse I-A/I-E Antibody (M5/114.15.2) (BioLegend, Cat # 107641, RRID:AB_2565975)
NK1.1: Brilliant Violet 785™ anti-mouse NK-1.1 Antibody (PK136) (BioLegend, Cat # 108749, RRID:AB_2564303)
SiglecF: BD Horizon™ BV421 Rat Anti-Mouse Siglec-F (E50-2440) (BD Biosciences, Cat # 562681, RRID:AB_2722581)
SiglecF: PE/Dazzle™ 594 anti-mouse CD170 (Siglec-F) Antibody (S17007L) (BioLegend, Cat # 155530, RRID:AB_2890716)
TCRβ: BUV737 Hamster Anti-Mouse TCR β Chain (H57-597) (BD Biosciences, Cat # 612821, RRID:AB_2870145)
TNF: APC anti-mouse TNF-α Antibody (BioLegend, Cat # 506308)
Live/Dead discrimination: LIVE/DEAD™ Fixable Aqua Dead Cell Stain Kit, for 405 nm excitation (ThermoFisher, Cat # L34966); LIVE/DEAD™ Fixable Blue Dead Cell Stain Kit, for UV excitation (ThermoFisher, Cat # L23105)
Tetramers: Anti-MHC class I TB10.4 tetramer (NIH Tetramer Core Facility, sequence: IMYNYPAM)

#### Cell sorting

Lungs were dissociated as described above and resuspended in RPMI (Gibco, #11875093) for labeling. Cell sorting was performed on a FACS Aria II (BD Biosciences). Sorted cells were collected in complete media, pelleted, resuspended in TRIzol, and frozen at -80°C overnight prior to RNA isolation.

### Confocal microscopy

Lungs were dissected and incubated in BD Cytofix diluted 1:3 with PBS for 24 hours at 4 °C. Lungs were then washed two times in PBS, incubated in 30% sucrose for 24 hours at 4 °C, embedded in OCT, and frozen in a dry ice slurry with 100% ethanol. 20 μm sections were cut using a CM1950 cryostat (Leica) and placed on charged slides. Sections were rehydrated with 0.1 M TRIS for 10 minutes at room temperature, incubated for 1 hour at room temperature with blocking buffer (0.1 M TRIS with 1% normal mouse serum, 1% bovine serum albumin, and 0.3% Triton X100), and then incubated overnight at room temperature with fluorescently conjugated antibodies or DNA dyes (Nucspot ® Nuclear Stains 750/780, Biotium, #41038; Mycobacterium tuberculosis purified protein derivative (PPD-Alexa488), Abcam, Cat # ab20962, RRID:AB_445945; Anti-mouse Histone H3Cit Abcam Cat # ab281584). Following labeling, slides were washed with 0.1 M TRIS for 30 minutes and sections sealed with coverslips and Fluoromount G mounting media (Southern Biotech, 0100-01). Images were acquired on a Leica Stellaris8 confocal microscope at room temperature using a 63X/NA1.20 HC PL APO water-coupled objective. For visual clarity, thresholds were applied to the displayed channel intensities using ImageJ with identical settings applied across experimental groups. To quantify the level of citrullinated histone H3 (Cit-H3) signal in each section, discrete lesions were identified visually based on purified protein derivative (PPD) antibody labeling and the fluorescent intensity of Cit-H3 labeling measured in 5 independent regions within the lesion. Background fluorescence was estimated using a similar analysis of unlabeled tissue sections.

### Gene expression analysis

#### Real-time PCR of lung tissue

The right superior lobe of the lungs was placed in TRIzol (Invitrogen, 15596018) and isolated using two sequential chloroform extractions, Glycoblue carrier (Invitrogen, AM9515), isopropanol precipitation, and washes with 75% ethanol. cDNA was synthesized using the RNA to cDNA EcoDry kit (Takara #693543) Expression of *Ifnb1* was measured using TaqMan primer probes (ThermoFisher, Mm00439552_s1), TaqMan Fast Universal PCR Master Mix (ThermoFisher, #4364103), and a Quant Studio 5 RT-qPCR detection system (ThermoFisher). Measurements were normalized to expression of *Eef1a1* expression in individual samples (Integrated DNA technologies - *Eef1a1* forward primer for custom TaqMan assay: 5’ GCAAAAACGACCCACCAATG 3’, *Eef1a1* reverse primer for custom TaqMan assay: 5’ GGCCTGGATGGTTCAGGATA 3’, *Eef1a1* probe for custom TaqMan assay: 5’/56-FAM/CACCTGAGCAGTGAAGCCAG/36-TAMSp/3’).

#### Bulk RNA-seq

RNA isolation was performed using TRIzol, two sequential chloroform extractions, Glycoblue carrier (Invitrogen, AM9515), 100% isopropanol precipitation, two washes with 70% ethanol, and final resuspension in RNase free water. RNA was quantified with the Bioanalyzer RNA 6000 Pico Kit (Agilent, 5067-1513). cDNA libraries were constructed using the SMARTer Stranded Total RNA - Pico Input Mammalian Kit (TaKaRa, 634411) following the manufacturer’s instructions. Libraries were amplified and then sequenced on an Illumina NovaSeq 6000 (150 bp paired-end). The read pairs were aligned to the mouse genome (mm10) using the gsnap aligner(*64*). Concordantly mapping read pairs (∼20 million / sample) that aligned uniquely were assigned to exons using the subRead program(*65*) and gene definitions from Ensembl Mus_Musculus GRCm38.78 coding and non-coding genes. Genes with low expression were filtered using the “filterByExpr” function in the edgeR package(*66*) from bioconductor.org. Differential expression was calculated using the “edgeR” package and false discovery rate computed with the Benjamini-Hochberg algorithm.

#### Single-cell RNAseq

Libraries were prepared using the Next GEM Single Cell 3′ Reagent Kits v3.1 (Dual Index) (10X Genomics, PN-1000268) following the manufacturer’s instructions. Raw sequencing data were aligned to the mouse genome (mm10) and UMI counts determined using the Cell Ranger pipeline (10X Genomics). Data processing, integration, and analysis was performed with Seurat v.3 (*67*). Droplets containing less than 200 detected genes, more than 4000 detected genes (doublet discrimination), or more than 5% mitochondrial reads were discarded. Genes expressed by less than 3 cells across all samples were removed. Unbiased annotation of clusters using the Immgen database (*68*) as a reference was performed with the “SingleR” package(*69*). Data visualization was performed with the “Seurat”, “tidyverse”, “cowplot”, and “viridis” R packages.

### Serum cholesterol analysis

Total cholesterol, HDL, LDL, and triglyceride levels were measured using the Rodent Lipid Panel by IDEXX BioAnalytics (Test Code 6290).

### Serum eicosanoid analysis

Mass spectrometry based lipidomic analysis was performed by the Cayman Chemical company. Prior to thawing the experimental samples, a mixture of the 19 calibration standards was prepared in methanol at a concentration of 270 ng/mL each. A series of nine 1/3 (v/v) dilutions was prepared in water/acetonitrile 1:1 (v/v), down to a concentration of 13.7 pg/mL. Fifty-microliter aliquots of these ten solutions were mixed with 100 µL of a methanolic solution containing 1 ng each of the internal standards and with 50 µL PBS to be processed for solid-phase extraction as described below. Quality control samples were also prepared independently by diluting a stock solution of calibration standards (1 µg/mL each) in water/acetonitrile 1:1 (v/v) to 200 ng/mL (HQC), 20 ng/mL (MHQC), 2 ng/mL (MLQC), and 0.2 ng/mL (LQC). Fifty-microliter aliquots of these solutions were mixed with 100 µL of a methanolic solution containing 1 ng each of the internal standards. After thawing, aliquots of the experimental samples analyzed (50 µL from serum) were transferred to a 96-well plate. To each sample, 100 µL methanol containing a mixture (1 ng each) of the internal standards was added, as well as 50 µL water/acetonitrile 1:1 (v:v). Samples were mixed well and placed at -80 °C overnight to improve extraction. They were then taken out of the freezer and thawed on wet ice, after which they were mixed thoroughly and centrifuged for 15 min at 770 x g. In the meantime, an appropriate number of wells on a 96-well solid-phase extraction (SPE) plate (Strata-X 33 µm Polymeric Reversed Phase, 10 mg, Phenomenex) were conditioned with 2 mL methanol and equilibrated with 2 mL water, using a nitrogen gas-driven positive-pressure manifold device from Biotage. All calibration and quality control samples, as well as the supernatants from the mouse serum samples, were transferred to a clean 2 mL 96-well plate and diluted to 900 µL with water. The plate was gently stirred, and samples were then transferred using a multichannel pipette onto the equilibrated SPE plate. After washing with 1 mL water and 1 mL water/methanol 9:1 (v/v), extracts were eluted with 0.9 mL methanol into a 96-well glass insert plate. Solvent was then evaporated using a SpeedVac concentrator, and the extract was resuspended in 100 µL water/acetonitrile 60:40 (v/v). Aliquots of 10 µL were injected into the LC-MS/MS system for analysis. The chromatographic profile of the ion count for each m/z transition was monitored, and the area under the peak (ion intensity vs elution time) integrated using commercial software (MultiQuant, Sciex). The area ratios of each analyte detected are interpolated in the calibration curve for the corresponding authentic standard, or in some cases for a structurally similar surrogate standard as listed on the accompanying data file. Calculations of the total amount of each oxylipin present in each sample were performed using MultiQuant. At least three quality control samples at each concentration level were run throughout the sample sequence to assess instrument performance, which was verified to be within an acceptable range throughout the sample queue. The full processed data set is available in the Supplemental material as Dataset S1.

### Study approval

All experiments were approved by the Institutional Animal Care and Use Committee at Seattle Children’s Research Institute and then performed in compliance with the relevant protocols.

### Statistical analysis

Statistical analysis was performed in R (v4.4.0). Definitions of center and dispersion are indicated in the figure captions. Measurements from individual replicates are indicated with points and unless otherwise noted indicate individual mice. Statistical significance was determined using the two-sided Student’s t-test allowing for unequal variances. Statistical significance of differences in measurements of bacterial burden by CFU analysis was assessed using the Wilcox rank-sum test. Significance of survival experiments was assessed using the log-rank (Mantel-Haenszel) test to test for a difference between Kaplan-Meier survival curves.

## Acknowledgments

We would like to thank the Office of Animal Care at the Center for Global Infectious Disease Research at Seattle Children’s Research Institute, for taking care of the mice. We would like to thank Dr. Sara Cohen, Dr. Courtney Plumlee, and Mari Morikawa for scientific advice and Nicholas Lee for assistance with data management.

## Author contributions

DL - Conceptualization, Investigation, Validation, Supervision, Writing (original manuscript)

DM - Investigation, Validation

ANJ - Investigation, Validation

TAM - Investigation, Validation JDA - Manuscript Review

BHG - Resources, Manuscript Review

KBU - Funding acquisition, Project Administration, Manuscript Review

AA - Funding acquisition, Project Administration

AHD - Conceptualization, Investigation, Validation, Software, Formal Analysis. Project Administration, Supervision, Writing (original manuscript)

ESG - Conceptualization, Investigation, Validation, Project Administration, Supervision, Writing (original manuscript)

