## Supplementary Figures for "APOE Protects Against Severe Infection with *Mycobacterium tuberculosis* by Restraining Production of Neutrophil Extracellular Traps"

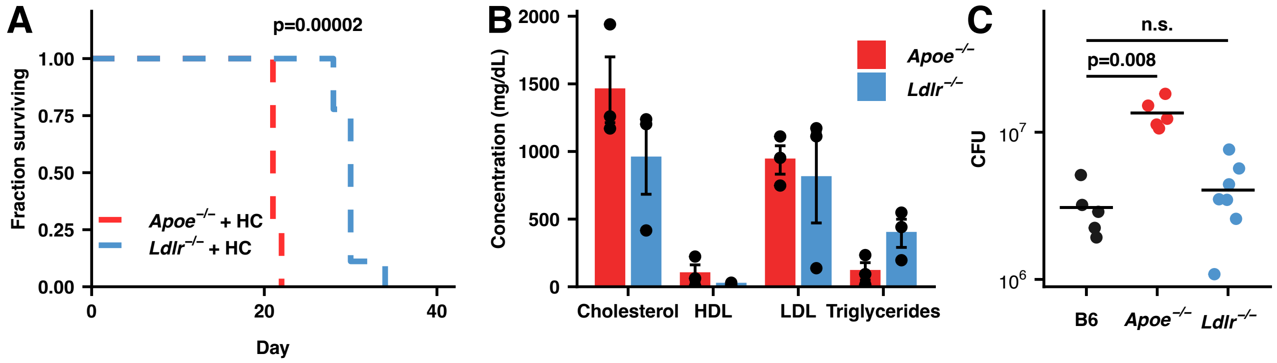


**Fig. S1.** ***Apoe**$^{-/-}$* **HC mice are highly susceptible to infection with Mtb**. (**A**) Female mice of the indicated genotypes were fed either normal food or high-cholesterol food for two weeks and then infected with ~50 CFU Mtb H37Rv and maintained on their pre-infection diet. (n=4-5 mice/group) (B) Serum cholesterol profiles at day 7 following infection of the indicated genotypes of mice fed HC food and infected with Mtb H37Rv as in (A). HDL = high-density lipoproteins, LDL = low-density lipoproteins. (n=3 mice/group) (**C**) Mice of the indicated genotypes were fed normal chow and then infected with ~50 CFU Mtb H37Rv. Bacterial burden in the lung was measured at day 28 PI by CFU counting. (n=5-7 mice/group) Bars/lines indicate mean; error bars indicate SEM. Significance analysis was performed using the Mantel-Haenszel test (A) or the Wilcox rank-sum test (C).

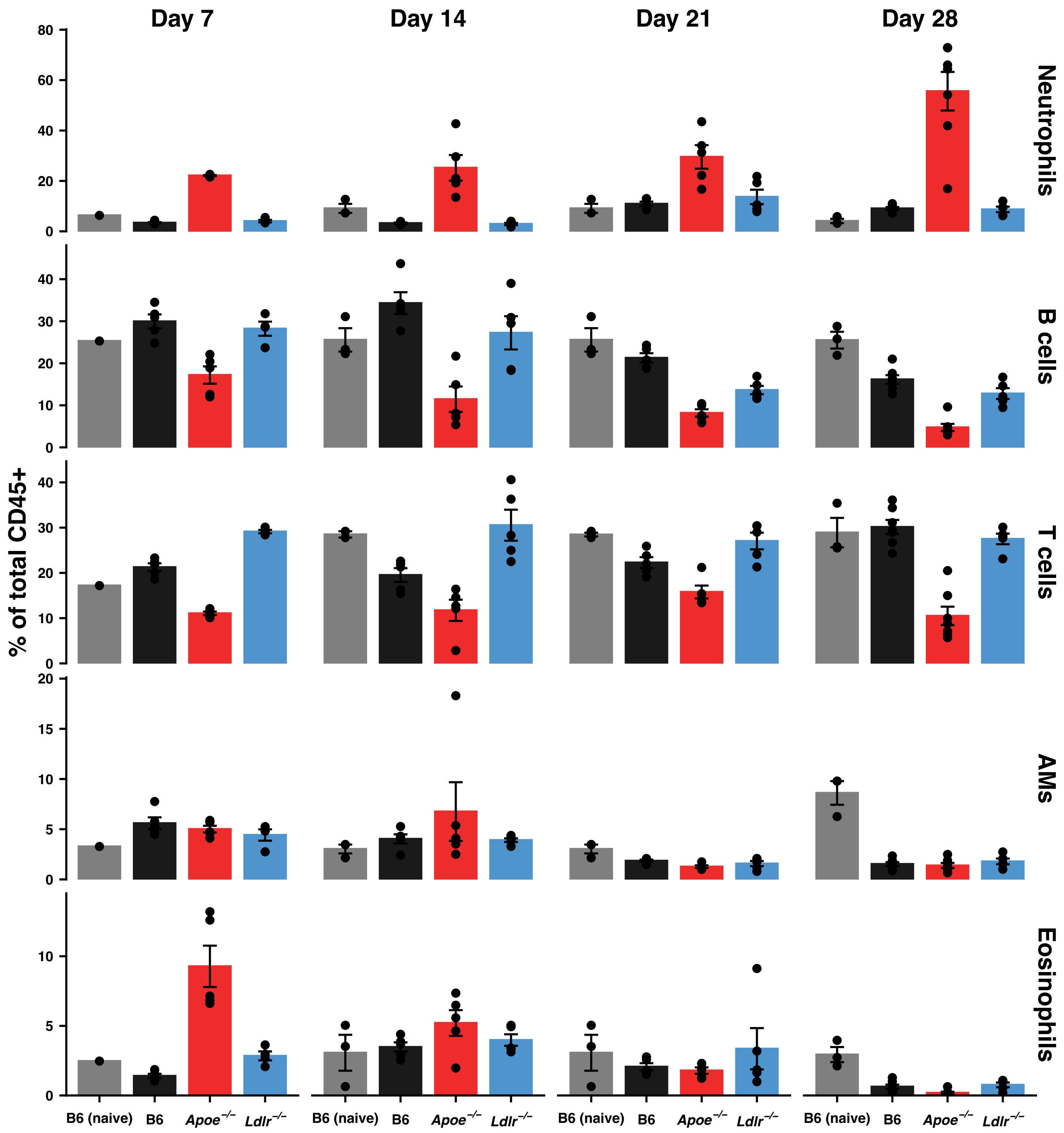


**Fig. S2. Pulmonary cellularity in B6, *Apoe*$^{-/-}$, and *Ldlr*$^{-/-}$ mice following Mtb infection.** Mice of the indicated genotypes on a HC diet were infected with ~50 Mtb H37Rv and the abundance of each cell type in the lung listed as a fraction of all CD45+ cells was determined by flow cytometry. Uninfected B6 mice on a normal diet were processed similarly for comparison. Bars indicate mean; error bars indicate SEM. See Figure S5 for gating strategy. (n=5-7 mice/group)

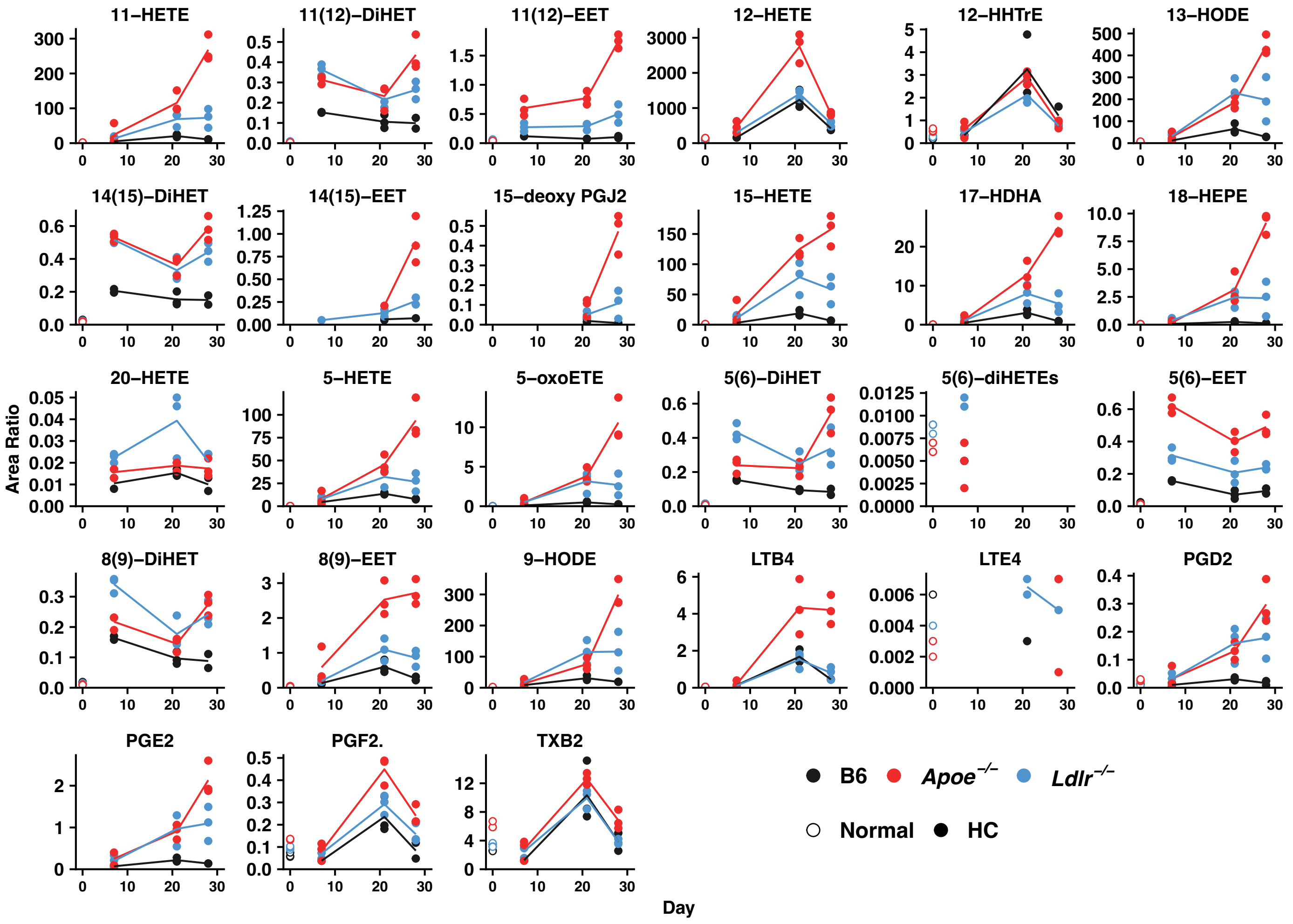


**Fig. S3. Serum eicosanoid levels in B6, *Apoe*$^{-/-}$, and *Ldlr*$^{-/-}$ mice following Mtb infection.** Serum was obtained at the indicated day PI and the abundance of each eicosanoid measured by mass spectrometry. The abundance of each species is calculated relative to an appropriate internal standard (See Methods). (n=3 mice/group)

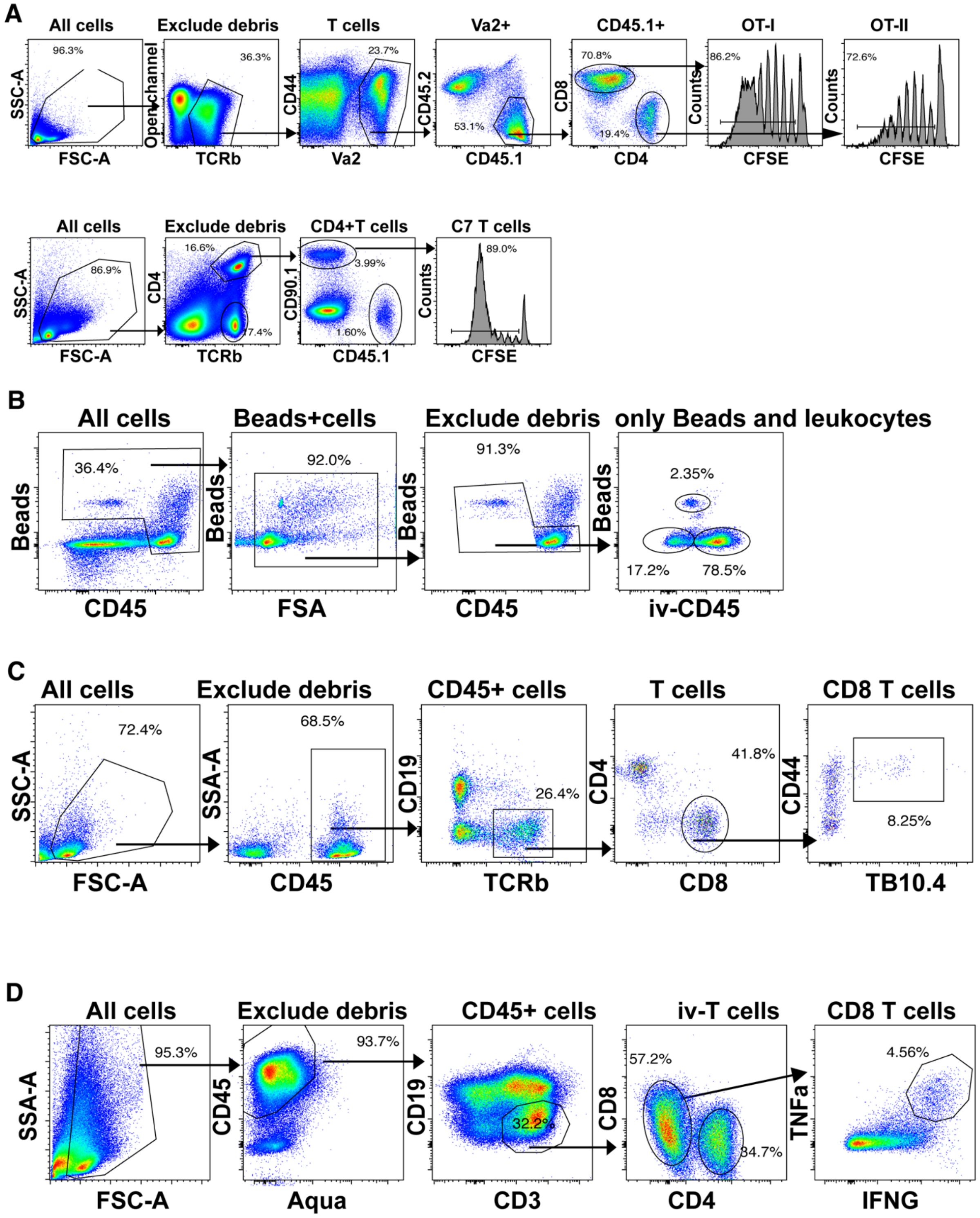


**Figure S4. Gating strategies for T cell priming experiments.** (**A**) *Expansion of adoptively transferred T cells in vivo:* The percentage of antigen-specific T cells dividing is computed relative to the total number of single T cells expressing CD45.1 (and CD8 or CD4 in the case of OT-II and OT-I T cells) (**B**) *Absolute counts of total T cells in the lung parenchyma:* Single CD45+ cells and counting beads are separated from debris by forward scatter and the absolute number of IV- cells computed relative to the known number of beads added. (**C**) *Fraction of antigen-specific T cells in the lung:* The percentage of TB10.4 tetramer+ T cells in the lung was defined relative to the number of single, CD45+ TCRb+ CD8+ CD44+ T cells. (D) *Ex vivo T cell restimulation:* The fraction of CD8 T cells producing both IFNG and TNF was computed relative to the total number of single, live CD3+ CD8+ T cells.

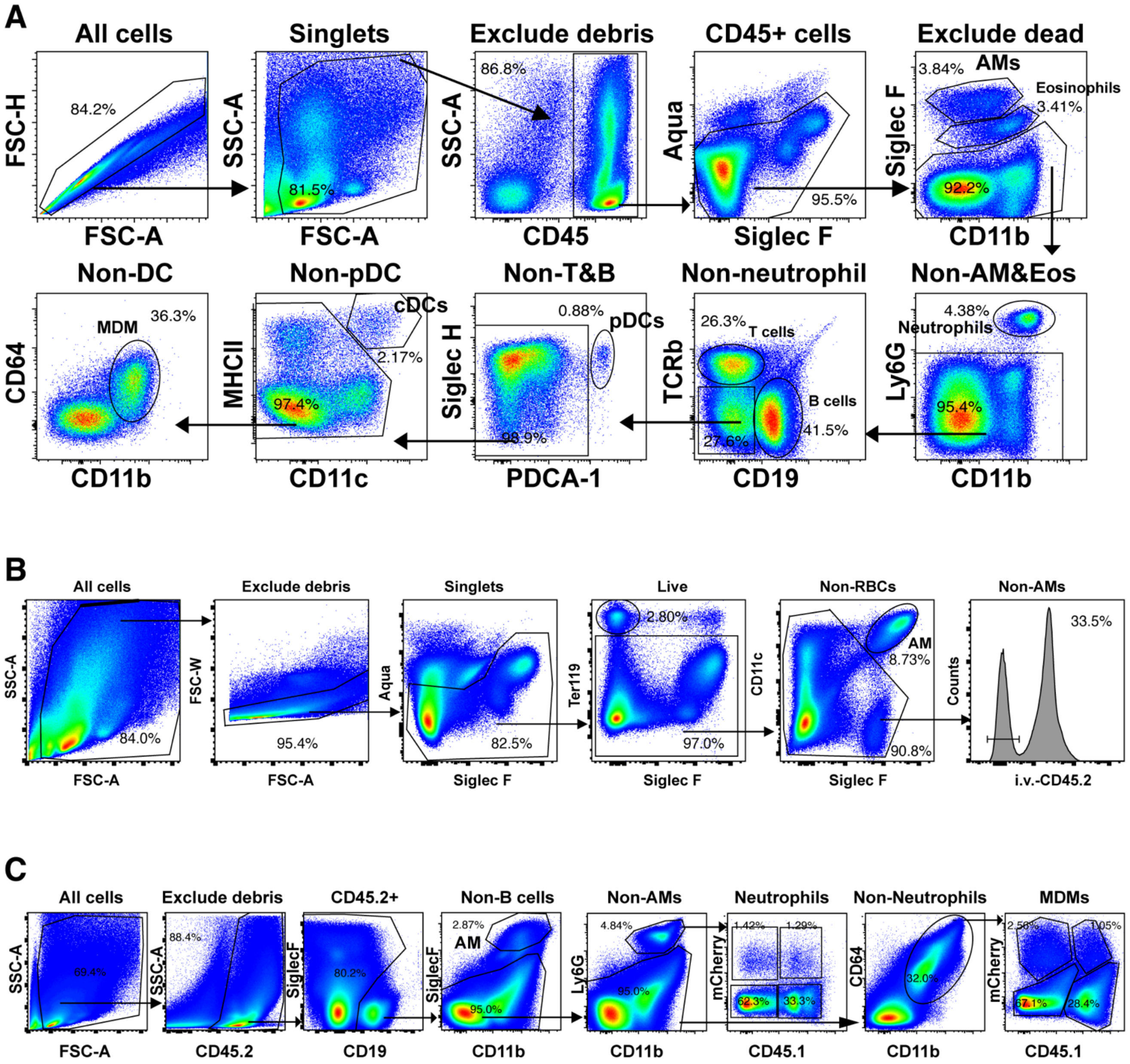


**Figure S5. Gating strategy for pulmonary cellularity, single-cell RNA-seq, and mixed bone-marrow chimera analysis.** (**A**) Single, live, CD45+ cells were identified as indicated. The fraction of CD45+ cells belonging to each of the indicated cell types was computed using the indicated gating hierarchy. (**B**) Prior to sacrifice, mice were injected with an PE-CD45.2 antibody to mark cells in the circulation. Following digestion of the lung to a single-cell suspension, single, live cells were gated on low Ter119 to exclude red blood cells and low PE-CD45.2 to define the population in the lung parenchyma. In addition, alveolar macrophages (AMs), were defined as SiglecF+ CD11c+ cells, regardless of PE-CD45.2 status (IV labeling is unreliable for AMs due to high background autofluorescence.). These two populations were combined and sorted for analysis by single-cell RNA-seq. (**C**) Mixed bone marrow chimeric mice on a B6 background (B6.SJL-Ptprc$^{a}$ Pepc$^{b}$/BoyJ (CD45.1)) were generated by reconstitution with a 50:50 mixture of B6 (CD45.1.2) and *Apoe*$^{-/-}$ (CD45.2) bone marrow. To isolate monocyte-derived macrophages (MDMs) and neutrophils single, CD45.2+, CD19- cells were selected. SiglecF+ alveolar macrophages (AMs) were excluded and infected/uninfected (mCherry+/-) Ly6G+ neutrophils of each genotype isolated by CD45.1 level. Infected/uninfected (mCherry+/-) Ly6G- SiglecF- CD64+ CD11b+ MDMs of each genotype were isolated by CD45.1 level.
